# EpiBlot: Joint Mapping of Chromatin Accessibility and Targeted Proteomics in HER2-expressing Breast Cancer Systems

**DOI:** 10.1101/2025.09.25.678463

**Authors:** Anna Fomitcheva Khartchenko, Trinh Lam, Nadine Goldhammer, Marin Bont, Shruti Warhadpande, Jennifer M. Rosenbluth, Amy E. Herr

**Affiliations:** University of California, Berkeley; University of California, San Francisco

## Abstract

HER2 proteoforms promote therapeutic resistance and aggressiveness in HER2-positive breast cancer, yet their epigenetic consequences remain poorly defined. Here, we establish EpiBlot, a joint assay incorporating a customized plateATAC-seq workflow that minimizes sample inputs with single-cell western blotting to concurrently profile chromatin accessibility with protein and proteoform expression. We applied our method to engineered MCF7 cells expressing HER2 proteoforms - full-length p185HER2 or truncated 611-CTF -, where we evaluated the impact of such proteoforms on the epigenetic and protein profiles after lapatinib or doxorubicin exposure. Expression of 611-CTF elicits pervasive chromatin remodeling, whereas p185HER2 provokes only modest accessibility shifts under the same treatments. EpiBlot reveals that treatment with doxorubicin drives extensive genome-wide accessibility changes, while lapatinib treatment produces limited global effects but unmasks proteoform-specific responses. Concordance between chromatin accessibility and protein abundance is moderate, underscoring complex regulatory coupling. Extending this dual-modality approach to HER2-low patient-derived organoids uncovers distinct chromatin states and reveals a subpopulation of triple-negative breast-cancer cells expressing truncated HER2 proteoforms. We anticipate that EpiBlot will highlight the value of multimodal profiling with proteoform identification for dissecting tumor heterogeneity and therapeutic response in cancer.

**Graphical abstract:** 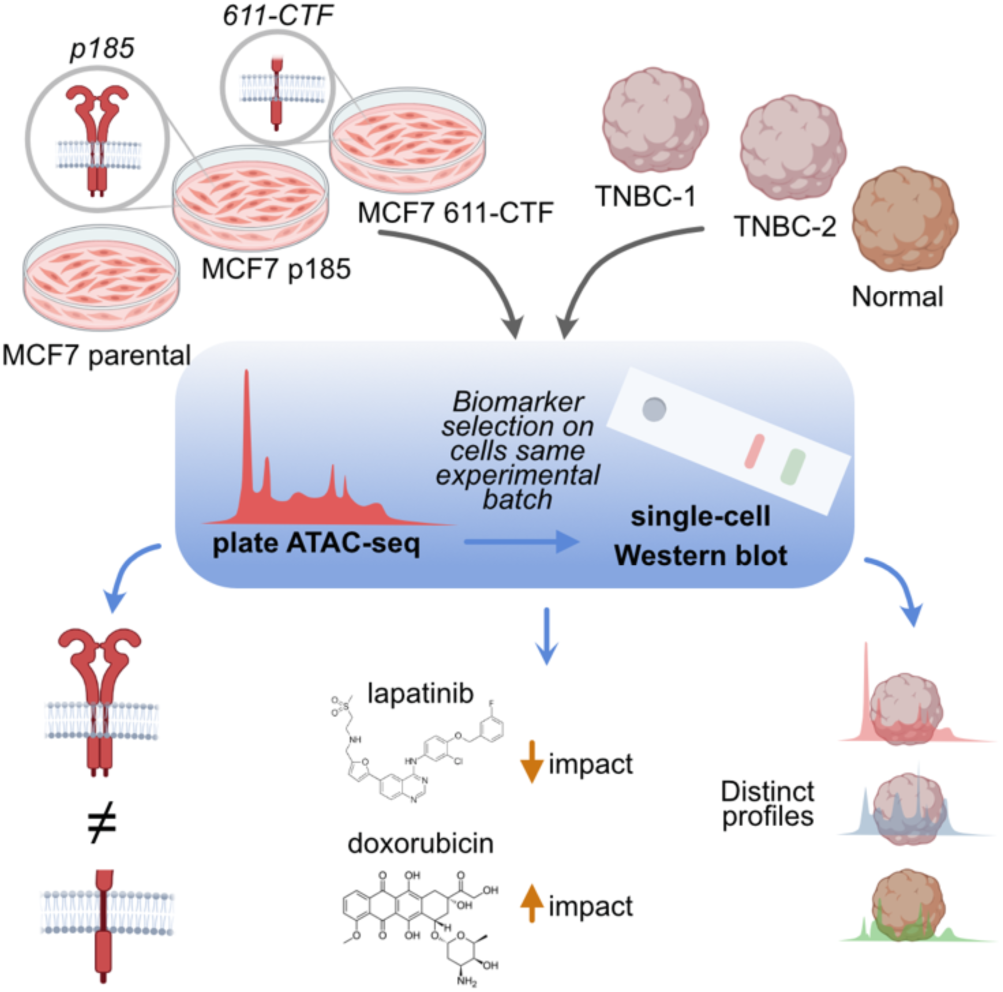

## Introduction

Female breast cancer was the second most commonly diagnosed cancer worldwide in 2022^1^. Among the subtypes of breast cancer, HER2+ accounts for up to 20% of all cases^2^. HER2 (human epidermal growth factor receptor 2) is a membrane-bound receptor that can form homodimers and heterodimers with other members of the Early Growth Response (EGR) family^3^. The signaling cascades downstream of HER2 regulate essential cell functions such as cell differentiation, proliferation, adhesion, and migration^3^. HER2 as a target for anti-cancer therapy remains a major focus of preclinical investigation^4^. Current anti-HER2 cancer therapies include monoclonal antibodies against the extracellular domain of HER2 (e.g., trastuzumab or pertuzumab) and small molecule inhibitors targeting the HER2 tyrosine kinase domain (e.g., lapatinib, neratinib, or tucatinib)^3^. While these treatments have been traditionally indicated for tumors with HER2 overexpression (HER2+ or HER2-amplified tumors), recent clinical advances show that patients with immunohistochemistry score of +1, or +2 with negative *in situ* hybridization for HER2, namely HER2-low or even HER2-ultralow tumors, can benefit from HER2-directed therapy, such as the antibody-drug conjugate trastuzumab-deruxtecan^5^. This emerging clinical category may represent a group with distinct biology that still remains under investigation^6,7^.

Multiple causes for therapeutic resistance in HER2+ cancers exist, including the expression of HER2 proteoforms. These proteoforms are altered forms of the full-length HER2 protein that include carboxy-terminus fragments (CTFs), splicing variants, and mutations. Select HER2 proteoforms have been explored for their impact in tumor progression and response to treatment. These are the d16HER2 splice variant, which results from the skipping of exon 16 during transcription, as well as 687-CTF and the 611-CTF proteoforms, produced by the alternative initiation of translation from the codon in the position 687 and 611, respectively^8^. These truncated forms are reported under the umbrella term p95HER2, which may include other truncated forms produced by the cleaving activity of metalloproteases, caspases, and calpains^8,9^. The p95HER2 proteoforms, present in approximately 30% of HER2+ patients, have been related to an increased aggressiveness, poor prognosis, and metastatic potential^8,10,11^. Notably, due to the absence of their extracellular domain, these proteoforms can evade therapies targeting the HER2 extracellular domain (i.e., trastuzumab). However, the p95HER2 proteoforms may still respond to combination strategies: trastuzumab paired with chemotherapy has shown efficacy, likely through chemotherapy-mediated sensitization^12^. Interestingly, HER2 levels in 611-CTF-expressing cells increased upon treatment with the chemotherapeutic drug doxorubicin^12^. The capacity of doxorubicin to induce de novo structural variants and genomic heterogeneity^13^ may impact the induction of HER2 expression in p95HER2+ cells. Although extracellular domain-targeting agents are ineffective against truncated HER2 proteoforms, intracellular inhibitors like lapatinib remain active. For instance, cells expressing p95HER2 treated with lapatinib inhibited cell growth due to a reduction of the downstream phosphorylation of Akt and mitogen-activated protein kinases (MAPK)^14^. These findings and recent initiatives in proteoform definition underscore the importance of proteoform profiling in guiding therapeutic decisions in diseases, including cancer^15^.

Integrating diverse data modalities has become instrumental in obtaining a complete picture of cellular states and unraveling tumor biology, since different molecular data types may result in varying degrees of biological redundancy. Protein data often shows an intermediate degree of redundancy with transcriptomic and epigenetic markers but has limited association to copy number aberrations^16^. Across proteins, it is now observed that proteoforms considerably expand the landscape of available protein forms, with many proteoforms potentially presenting diverse functions in the cell^17^. Epigenetic markers are increasingly recognized as key determinants in tumor progression, metastasis, and tumor fate^18,19^. Among these epigenetic markers, chromatin state shifts have been observed across basal and luminal breast cancer lineages, within molecular subtypes, and between primary tumors and metastatic sites^20–23^. Epigenetic modifications frequently precede changes in protein expression, with chromatin structure predictive of coordinated RNA-protein regulatory networks^22^. Nonetheless, the correlation between RNA and protein is only intermediate^16^, reinforcing the value of integrating multiple data modalities, even among moderately redundant ones, to comprehend and predict tumor behavior^16,24,25^. For this reason, in studying the role of HER2 proteoforms, we aim to explore the influence of such proteoforms on epigenetic regulation by using chromatin accessibility and protein expression as our measurement types. Although multiple methods exist for simultaneous chromatin accessibility and protein analysis, such as bulk approaches combining chromatin accessibility^26^ analysis with proteomic techniques, or single-cell methods such as TEA-seq^27^, DOGMA-seq^28^, or NEAT-seq^29^, to the best of our knowledge, a key limitation is the detection of proteoforms.

To address this gap, we introduce EpiBlot (**Figure 1A**), an integrated multiomic analysis of HER2- related breast cancer model systems by combining chromatin profiling by transposase-accessible chromatin with sequencing (ATAC-seq)^26^ and targeted protein expression analysis by single-cell western blotting (scWB)^30–33^. To perform ATAC-seq, we developed a plate-based method utilizing a mini-96-well plate that accommodates less than 20 µL total reaction volume and requires only 5–500 nuclei, without the need for a thermoshaker during the tagmentation step. The scWB analyzes protein and proteoform expression with single-cell resolution. One of the main advantages that lead us to choose the scWB as a counterpart to ATAC-seq is the capability of the scWB to stably archive solubilized protein targets by covalent bonding to the gels^34^. This single-cell lysate archiving allows retrospective protein analysis. Thus, we selected our protein targets based on the differential chromatin accessibility observed in the ATAC-seq dataset. While ATAC-seq was used for target selection, the use of a single-cell technique for protein analysis explores intratumoral heterogeneity landscapes.

**Figure 1.**
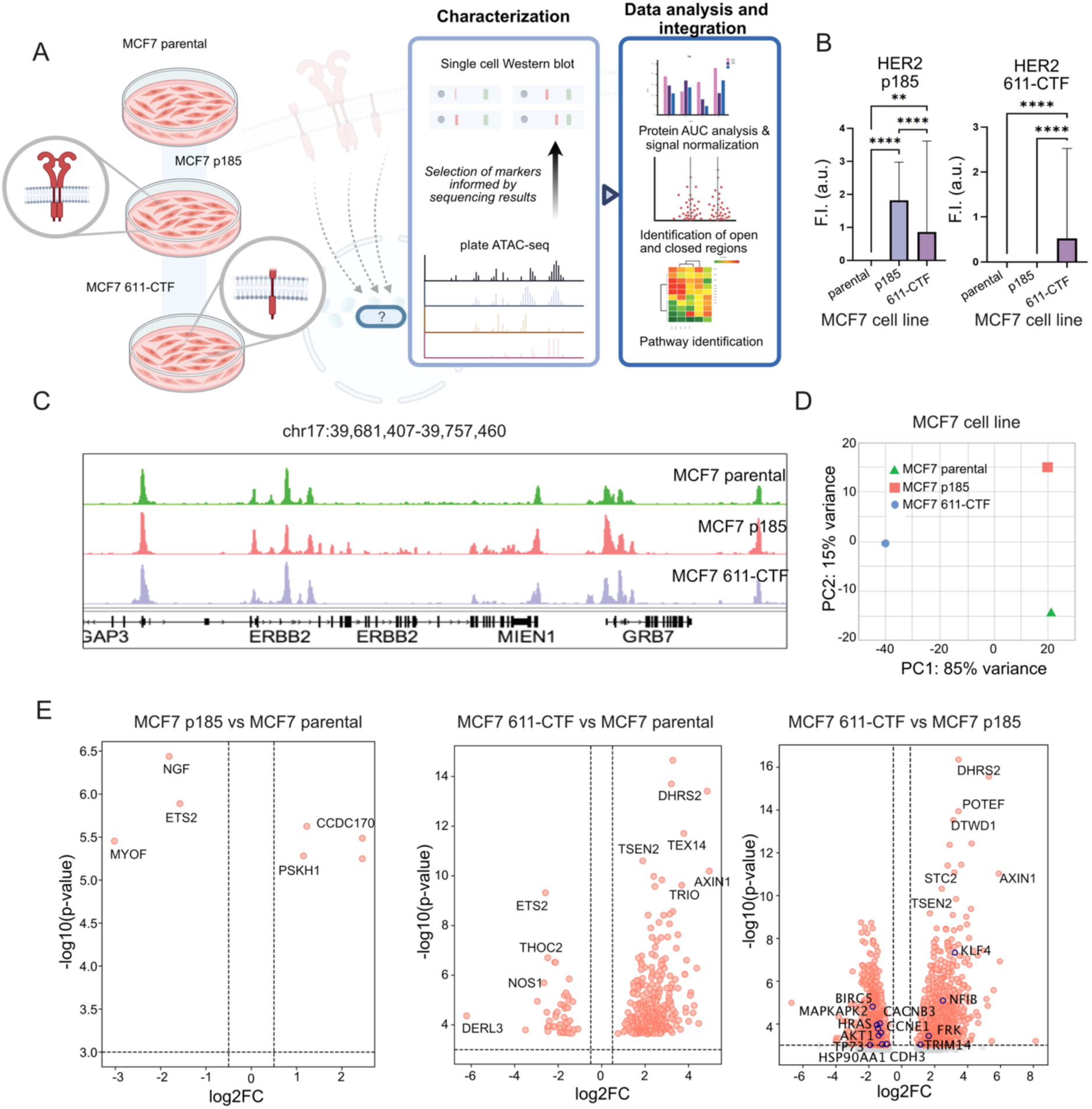
The presence of HER2 611-CTF alters chromatin accessibility more substantially than increased abundance of p185 proteoform. A) Schematic of the study. Three transfected cell lines with HER2 proteoforms (empty vector - parental-, full-length HER2 form p185, and truncated proteoform 611-CTF) are evaluated for the changes in chromatin accessibility, and protein expression based on the selected genes from the sequencing experiment. . B) Bar plot showing HER2 p185 (left) and 611-CTF (right) expression across the transfected cell lines, confirming that MCF7 p185 present only p185 expression, while MCF7 611-CTF have expression of both full-length and fragment protein. F.I. corresponds to fluorescence intensity and a.u. to arbitrary units. C) Changes in chromatin accessibility at the ERBB2 gene in MCF7 parental, p185, and 611-CTF cell lines. Genome browser tracks (GRCh38) show chromatin accessibility at the ERBB2 (HER2), MIEN1, and GRB7 gene loci in MCF7-transfected cell lines. Differentially accessible (DA) peaks are enriched in the ERBB2 region in the full-length HER2 (p185) sample, while DA peaks in GRB7 are observed in both p185 and 611-CTF HER2-expressing cell lines. All track lines have the same y-axis limits, and peak heights are scaled across all samples. D) Principal Component Analysis (PCA) of ATAC-seq signal in the three MCF7-transfected cell lines. The PCA plot shows sample clustering based on chromatin accessibility variation. PC1 (85% variance) separates p185 (red) and parental MCF7 (green), while 611-CTF (blue) clusters separately from both, indicating a distinct chromatin accessibility profile compared to the other conditions. E) Volcano plots of ATAC-seq peaks comparing p185 vs. parental MCF7s (left), 611-CTF vs. parental MCF7 (middle), and 611-CTF vs. p185 (right). Peaks with significantly differential chromatin accessibility are highlighted in red, while the non-significant peaks are grey. The vertical dotted line represents an absolute log2 fold change of 1.0, while the horizontal dotted line represents an adjusted p-value threshold of 0.00001. FDR-corrected p-values were obtained using DESeq2.

Using EpiBlot, we successfully profiled chromatin accessibility and protein expression in MCF7 cells transfected with different HER2 proteoforms for their multiomic profiles, revealing that HER2 611-CTF broadly impacts chromatin expression. We then analyzed how the presence of HER2 proteoforms affects the response to two different drugs: a chemotherapeutic agent (doxorubicin) and a HER2-targeted tyrosine kinase inhibitor (lapatinib). While doxorubicin produced a widespread effect across the chromatin landscape, lapatinib induced only moderate changes. Finally, we extended our analysis to clinical samples, with three patient-derived breast cancer organoids (PDOs) – two triple negative breast cancers (TNBCs) classified as HER2-low and one from normal tissue as a control – to evaluate the chromatin profiles and HER2 proteoform expression.

## Material and methods

### Transfected MCF7 cells

Transfected MCF7 cell lines were kindly provided by the Arribas lab, with the cells reported and characterized in ref.^9^. MCF7 cells were cultured in DMEM/F-12 (10565018, Thermo Fisher), Glutamax (35-050-061, ThermoFisher) supplemented with 10% FBS (S11150, Gemini Bio), 0.2 mg/mL Geneticin G418 (10131035, Thermo Fisher) and 1 μg/mL doxycycline (63131, Takara Bio) at 37°C, 5% CO2. Media was exchanged every 48 h to avoid activation of the vector. MCF7 were maintained by dissociation into single cells at 70-80% confluency with Trypsin-EDTA 0.25% and seeded at a dilution of 1:10 in media. For expression of the TET-off vector, the cells were cultured in media without doxycycline for 48 h.

For the drug treatment, cells were cultured in doxycycline free media for 48 h. After, 1 μM of doxorubicin (C819Z82, Thomas Scientific) or 10 μM of lapatinib (SIAL-CDS022971-25MG, Sigma-Aldrich) were deposited in the corresponding wells of the 96-well plate. For the controls, a volume of DMSO (12611S, Cell Signaling Technology) equivalent to that of doxorubicin or lapatinib in the media was added to the cells to account for the toxicity of the DMSO, the initial diluent of these drugs.

To reduce the impact of batch variation, all samples, including all treated cells with doxorubicin and lapatinib, were processed in parallel.

### Organoid culture

PDOs from the Rosenbluth lab organoid biobank were cultured as described previously^35–37^. **Supplementary Table 1** catalogues the PDOs scrutinized here. Briefly, PDOs were embedded in Cultrex Basement Membrane Extract (BME, 3532-010-02, BioTechne) type 2 and plated in preheated (37°C) 24-well non-cell treated plates. TNBC PDOs were kept in type 1 organoid medium, while normal PDOs were kept in type 2 organoid medium. Organoid medium is composed of Advanced DMEM/F12 (BW12-719F, Fisher Scientific) with 50 μg/ml Primocin (ant-pm-05, InvivoGen), 1x GlutaMax, 100 units/ml Penicillin/Stroptomycin (15140122, Thermo Fisher), 10 mM HEPES (15-630-080, Thermo Fisher), 20% Wnt3a conditioned medium^37^ (H. Clevers lab, Hubrecht Institute, type 2 only), 10% R-spondin 1 conditioned medium^37^ (C. Kuo lab, Stanford University), 10% Noggin conditioned medium^37^ (H. Clevers lab, Hubrecht Institute), 1x B27 (17504044, Gibco), 10 μM Nicotinamide (N0636, Sigma-Aldrich), 1.25 mM N-acetylcysteine (A0737, Sigma-Aldrich), 500 ng/ml Hydrocortisone (type 2 medium only, H0888 Sigma-Aldrich), 100 nM β-estradiol (type 2 medium only, E4389 Sigma-Aldrich), 10 μM forskolin (type 2 medium only, F6886 Sigma-Aldrich), 5 μM Y-27632 (ab143784, abcam), 5 μM heregulin β1 (100-03, Peprotech), 5 μM FGF-7 (type 1 medium only, 100-19 Peprotech), 20 ng/ml FGF-10 (100-26, Peprotech), 500 nM A83-01 (SML0788, Sigma-Aldrich), 5 ng/ml EGF (AF-100-15, Peprotech), and 1 μM SB202190 (type 1 medium only, S7067 Sigma-Aldrich). To obtain single cells for analysis, organoid domes were digested for 10 min with TrypLE Express (12604013, Gibco) at 37°C followed by three washes with Advanced DMEM/F12 with 50 μg/ml Primocin, 1x GlutaMax, 100 units/ml Penicillin/Streptomycin, and 10 mM HEPES. Single cells were harvested in DPBS (14190144, Gibco) with 5% FBS (SH30910.03, Cytiva) and kept on ice until analysis. For the tests, Normal had 1 replicate, TNBC-1 had 2 replicates, and TNBC-3 had 3 replicates.

### Immunofluorescent staining of cell lines

Coverslips were placed at the bottom of a sterile 12-well plate and sterilized by washing in 70% ethanol 3x for 5 min. A fourth wash was performed with 100% ethanol for 5 min and the coverslips were dried inside the hood. A solution of poly-lysine was prepared by diluting 5 mg of poly-lysine in 50 mL of sterile water. The coverslips were coated with poly-lysine by submerging in 0.5 mL of the poly-lysine solution for 5 min. After, the coverslips were rinsed with water and dried for 2 h.

Cells were trypsinized and cultured on the coverslip inside a 12-well plate, following the culturing conditions described in “Transfected MCF7 cells”. After, the media was removed and cells were washed in PBS (10010023, Thermo Fisher). The PBS was substituted by 4% paraformaldehyde (28908, ThermoFisher) diluted in PBS for 20 min. The coverslips were washed 2x with PBS and the cells were permeabilized with 0.25% Triton X-100 in PBS for 5 min. Cells were then washed 2x in PBS.

The next step was the primary antibody staining. A solution of 1% BSA in PBS solution with 2.5 μg/mL of anti-HER2 antibody (ab16901 [discontinued], ab134182 (**Supplementary** Figure 1), abcam), 1 μg/mL of Hoescht 33342 (H1399, Invitrogen), and 0.033 μM of phalloidin-rhodamine (R415, Invitrogen) was prepared. A volume of 125 μL of the antibody solution was added to each coverslip and then incubated at 37°C for 3 h in a humidity chamber. The coverslips were then washed 5 times for 3 min in 0.1% Tween-20 in PBS on a rotator. For the secondary antibody staining, a solution containing anti-mouse-488 antibody at 2.5 μg/mL in 1% BSA with PBS was prepared and the coverslips were incubated in 125 μL of antibody solution for 1 h at 37°C in a humidity chamber. The coverslips were then washed in 0.1% Tween-20 in PBS 5x for 3 min. A drop of mounting media was deposited on the coverslip, and a glass slide was sealed on top. The slide was dried overnight and sealed with nail polish before imaging using the Agilent/Biotek LionHeart system.

### Immunofluorescent staining of PDOs

PDOs were digested to single cells as described above and fixed in 2% paraformaldehyde for 15 min. Cells were washed 3x with PBS and loaded onto a double cytology funnel (10-356, Fisher Scientific). Cells were spun onto microscope slides by centrifugation at 800 rpm for 3 min using a Thermo Scientific Cytospin 4 Centrifuge. Slides were then air-dried for 1 h followed by permeabilization with cold 0.5% Triton-X in PBS for 10 min at 4 °C. Subsequently, slides were washed 3x with PBS and then blocked with 10% goat serum in TBS-T for 1 h at room temperature. Slides were incubated overnight with an anti-HER2 antibody conjugated to AF488 (1:200) (ab275994, abcam) diluted in 1% goat serum in TBS-T. After 3x washes with TBS-T, slides were mounted with ProLong™ Gold Antifade Mountant with DAPI (P36931, Invitrogen) and imaged using an ECHO Revolve Fluorescent Microscope.

### Isolation of nuclei

ATAC-RSB (10 mM Tris-HCl pH 7.4, 10 mM NaCl, 3 mM MgCl₂) was prepared from stock reagents and supplemented to make lysis buffer (0.1 % IGEPAL CA-630, 0.1 % Tween-20, 0.01 % digitonin (G9441, Promega)) or wash buffer (0.1 % Tween-20). One million viable cells were pelleted (500 × g, 5 min, 4 °C), washed once in ice-cold PBS, and lysed in 300 µL lysis buffer (10 gentle mixes, 5 min on ice; 4 min for organoids). Lysis was quenched with 1 mL wash buffer, nuclei were pelleted 2x (hinge in/out, 500 × g, 3 min, 4 °C), and resuspended in 250 µL wash buffer with a wide-bore tip. Nuclei counts were determined on a Countess (Trypan Blue). Aliquots of 100,000 nuclei were diluted, re-pelleted, and adjusted to the desired concentration.

### Plate-based tagmentation and library preparation

PlateATAC-seq begins by tagmenting ∼300 nuclei in a single 15 µL microtiter plate microwell. A tagmentation mix was prepared using 1X Tagmentation buffer (20034197, Illumina Inc), 0.1% Tween 20, and 0.01% digitonin (Promega). TDE1 tagmentation enzyme (Tn5, Illumina Inc) was added to the mix at a ratio of 1:15. Then, 1.5 µL of nuclei (∼300 nuclei) were aliquoted into a mini- 96-well plate (Kubo Biotech), followed by the addition of 2.5 µL of the tagmentation mix. The plate was sealed with PCR tape and incubated for 30 minutes at 37°C in a qPCR instrument (q225, Kubo Biotech).

Instead of stopping the reaction with an elution-and-column cleanup, as in conventional bulk ATAC-seq^26^, we quench Tn5 by adding EDTA to chelate MgCl₂ directly in the micro-titer plate microwell. To terminate the tagmentation reaction, 2 µL of stop buffer (0.025 M EDTA, 0.015 M Tris-HCl, pH 8.0) was added to each microtiter plate microwell, and the mixture was incubated for 30 minutes at 50°C. For library amplification, 9 uL of PCR mix containing 1X NEBNext® High-Fidelity 2X PCR Master Mix (M0541S, NEB), 1.25 µM i5 primer, and 1.25 µM i7 primer was added to each microtiter plate microwell. The amplification protocol was as follows: 72°C for 5 min and 98°C for 30 s; followed by 10 cycles of 98°C for 10 s, 63°C for 30 s, and 72°C for 60 s. PCR products were purified using the Zymo DNA Clean and Concentrator-5 kit. DNA concentration was quantified using a Qubit fluorometer, and the nucleosomal pattern was assessed using an Agilent TapeStation with D1000 tape and reagents.

Libraries from all MCF7 cell lines or organoids were quantified with the KAPA Library Quantification Kit (07960140001, Roche), pooled equimolarly to 1.07 ng µL⁻¹ for cell lines and 1.70 ng/uL for organoids, and quality-checked by TapeStation and Qubit. The pooled library containing the cell lines was sequenced on two NovaSeq 6000 S2 lanes (∼3 billion clusters total) with 2 × 75 bp reads while the organoids pooled library was sequenced on NextSeq 2000 P2 (400 million clusters).

### Soft lithography for fabrication of single-cell western blot microdevices

Soft lithography for scWB molds were fabricated as previously reported^30^. A 40-μm layer of SU- 8 3050 was spun over a silicon wafer for 10 s at 500 rpm with an acceleration of 100 rpm, and then for 30 s at 4000 rpm with an acceleration of 300 rpm. The wafer with the SU-8 later was soft baked for 2 min at 65°C, 15 min at 95°C and then 3 min at 65°C. The wafer was then exposed to a UV dose of 385 mJ/cm2 and baked post-exposure for 1 min 65°C, 5 min at 95°C and then 1 min at 65°C. To remove excess of SU-8, the wafer was developed for 5 min on a shaker and then hard baked at 200°C for 20 min.

### Single-cell western blotting

The scWB was performed as previously described^30,38^. Briefly, gels were fabricated mixing 7% acrylamide:bis-acrylamide (29:1, A3574 Sigma-Aldrich), 3 mM BPMA (Raybow Pharmaceutical, custom), 0.08% ammonium persulfate (A3678-25G, Sigma-Aldrich), and 0.08% tetramethylethylenediamine (T9281-25ML, Sigma-Aldrich). The precursor solution was introduced to the gap between a patterned wafer and a silanized glass slide until the gel was polymerized after 15 min. The gel was detached from the wafer and washed in PBS for 1h. The gel was then dried under a nitrogen stream to increase the probability of deposition of a cell in each microwell.

For cell lines, 250 μL of 1 million/mL of strained cells were deposited on each gel. The dimensions of the gel correspond to half of a microscope glass slide. In the case of organoids, in all cases the concentration of cells was below 1 million/mL and thus the available concentration was deposited on the gel in 250 μL. After 10 min, the excess cells were carefully removed by tilting the slide and washing it with PBS. The gels were deposited in a custom 3D printed chamber that contained two platinum electrodes separated by 6 cm. The cells were lysed by applying RIPA buffer (1% w/v SDS (SIAL-L3771-1KG, Sigma-Aldrich), 0.5% w/v sodium deoxycholate (30970, Sigma-Aldrich), 0.1% v/v Triton X-100 (SIAL-X100-100ML, Sigma-Aldrich), and 0.5× of Tris-glycine (93321-1L, Sigma-Aldrich)) for 30 s and then energizing an electrical field of 40 V/cm was applied for 30 s to perform electrophoresis. The gels were photoactivated (45 s) to covalently immobilize proteins with the BPMA in the polyacrylamide gel. The gel was then deposited in a volume of 1× TBS + 0.1% Tween-20 (TBST) for >1 h. The gel was then incubated in 40 μL of primary antibody for 1 h (100 μg/mL for β-tubulin (ab6046, abcam), TFAP2C (ab134182, abcam), GATA3 (14- 9966-82, ThermoFisher), and HER2, 42.5 μg/mL for BCAS3 (ab71162, abcam), 20 μg/mL for PPP1R13L (PA5-59491, ThermoFisher), 10 μg/mL for ULK1 (8054S, TheremoFisher), 9 μg/mL for EGR1 (MA5-15008, ThermoFisher)), washed in TBST for 45 min with one wash exchange after 20 min, and incubated with the secondary antibody for 1 h (100 μg/mL). The sample was washed in TBST for 45 min with one wash exchange, and then rinsed in water to remove salts. The gel was dried under a nitrogen stream and imaged using Genepix Microarray Scanner (Genepix 4300A, Molecular Devices), with a resolution of 5 μm.

### Data analysis

All ATAC-seq downstream data analyses were performed in R 4.2.2. Upstream data analyses including peak calling and per-sample quantification (featureCounts) were produced by the nf-core/atacseq v2.0 pipeline, then imported as a DESeqDataSet. Sample metadata were parsed from file names to create a design ∼group, and differential accessibility was assessed with DESeq2 (Wald test, BH < 0.05). Peaks were annotated to nearest genes (±3 kb TSS) with ChIPseeker / TxDb.Hsapiens.UCSC.hg38.knownGene, functional enrichment was performed with clusterProfiler::enrichGO, and results were visualized with ggplot2, pheatmap, and EnhancedVolcano.

scWB protein data analyses were performed using a custom the Python algorithm workflow adapted from ref.^39^, using numpy, pandas, tkinter, and pillow (**Supplementary** Figure 2, link: https://github.com/herrlabucb/segmentation_scWB). All analyses of scWB were performed on raw image data. For improved visualization, figures showing the scWB protein bands have been grayscale inverted and adjusted for contrast.

Nuclear characteristics (circularity, length and area) were quantified using MicrobeJ^40^ in Fiji v1.54d. The segmentation of the nucleus and the cytoplasm for all conditions was performed using UNSEG^41^. Schemes were created with BioRender.

### Statistical tests

Bar graphs were compared using two-way ANOVA followed by Tuckey’s multiple comparison tests. Nuclear dimensions were compared using the Mann–Whitney U test to account for skewed data using GraphPad Prism.

## Results

### Validation of EpiBlot on HER2-proteoform expressing cell lines

As a first step, we verified that differences across proteoform-expressing cell lines could be detected by both the scWB and plateATAC-seq independently. We evaluated TET-off MCF7 cell lines engineered to express distinct HER2 proteoforms, kindly provided by the Arribas lab^9^. These cell lines provide three conditions: MCF7 parental, transduced with an empty vector; MCF7 p185, containing the full-length HER2 protein (185 kDa); and MCF7 611-CTF, expressing the truncated 611-CTF HER2 fragment (110 kDa). We validated the expression of the HER2 protein using a scWB (**Figure 1B**). As expected, MCF7 parental cells presented minimal HER2 signal, consistent with null to low HER2 expression in MCF7 cells^42^. MCF7 p185 presented only the expected full-length proteoform (185 kDa). In contrast, MCF7 611-CTF presented both the full-length and the truncated proteoforms. Immunofluorescence staining confirmed HER2 localization to the cell membrane (**Supplementary** Figure 3). We also observed changes in the nuclear structure in each cell line, observing significant changes in the area (**Supplementary** Figure 4**, Note 1**).

To assess the impact of HER2 proteoforms on chromatin organization, we compared the chromatin accessibility profile across the three engineered MCF7 cell lines. To do so, we introduced a modified ATAC-seq protocol, called plateATAC-seq. The plateATAC-seq tagments ∼300 nuclei, carrying all the PCR steps sequentially in that same microtiter plate microwell, eliminating the tube transfer and column cleanup post-tagmentation, minimizing sample loss, and dispensing with the thermal shaker required to keep 50,000-nucleus bulk reactions in suspension. Because all steps remain on-plate, the workflow is inherently automation-friendly, allowing microtiter plate microwells to be processed in parallel and thereby maximizing experimental scale while conserving precious nuclei (**Supplementary** Figure 5A).

We first optimized the plateATAC-seq workflow by titrating the Mg²⁺-chelating quench with EDTA and the amplification parameters (**Supplementary** Figure 5B). Stopping tagmentation with 25–50 mM EDTA at 50 °C, followed by 12 PCR cycles, maximized library yield while preserving fragment integrity; the cycle number can be proportionally reduced for larger nuclear inputs. Using these conditions, we generated 43 libraries from MCF7 cells subjected to five treatment conditions. **Supplementary** Figure 5C presents the quality of the pooled library of 43 samples with TapeStation electropherogram displaying well-resolved nucleosomal peaks: nucleosome-free (∼180–200 bp), mononucleosome (∼349 bp), and dinucleosome (∼550 bp), confirming proper fragment sizing. In parallel, qPCR amplification curves for three serial dilutions exhibit the expected log-linear shift, further validating library integrity post-pooling. Four standard QC metrics are summarized in **Supplementary** Figure 6. The plateATAC-seq protocol consistently achieved a median fraction of reads in peaks (FriP) of ∼0.60, library complexities of 3–4 × 10⁷ unique fragments, and mitochondrial read fractions below 0.20. Taken together, these are values that match, and in some cases exceed, those reported for conventional bulk (50,000- nucleus) or commercial single-cell ATAC-seq platforms^43^ as shown in **Supplementary** Figure 6A. Genome-browser tracks confirmed that chromatin-accessibility landscapes generated from 300-nucleus plateATAC-seq libraries recapitulate those obtained from 50,000-nucleus bulk preparations at both genome-wide and locus-specific levels (**Supplementary** Figure 6B). Insert-size distributions retained the expected mono- and dinucleosomal periodicity (**Supplementary** Figure 6C), and transcription-start-site metaplots showed the characteristic peak at 0 bp with symmetrical decay across ± 3 kb (**Supplementary** Figure 6D). Together, characterization of plateATAC-seq shows that performing tagmentation, quenching, and PCR sequentially in a single 15 µL microtiter plate microwell not only preserves assay fidelity but also obviates tube transfers and column clean-ups, making the method readily automatable and ideally suited to precious or limited samples – including the PDO models profiled here.

We then profiled chromatin accessibility across our three MCF7 models: the parental line, cells expressing full-length HER2 (p185), and cells expressing the truncated 611-CTF proteoform, to identify proteoform-specific regulatory signatures. A representative locus is shown in **Figure 1C**: accessibility tracks at the *ERBB2* (HER2) gene reveal distinct peak architectures among the three lines, illustrating the pronounced remodeling driven by the 611-CTF isoform relative to both the parental and p185 cells. Genomic feature distributions were comparable across cell lines (**Supplementary** Figure 7A). Following peak calling from ATAC-seq data across the genome and gene-associated regions, we performed principal component analysis (PCA), which revealed that MCF7 parental, p185, and 611-CTF cells exhibit distinct chromatin accessibility profiles. The first two components accounted for 85% and 15% of total variance, respectively (**Figure 1D**). To further quantify these differences, we performed differential accessibility analysis using DESeq2. This analysis identified significantly differentially accessible regions (DARs) (adjusted *p* < 0.05, |log₂FC| > 1), as shown in the volcano plots (**Figure 1E**). MCF7 parental and MCF7 p185 presented minimal differences in the differentially accessible chromatin regions, consistent with their shared expression of HER2, despite different abundance levels. The 611-CTF fragment, however, led to substantial changes in chromatin accessibility, resulting in distinctly differentially accessible chromatin regions to both MCF7 parental and MCF7 p185 cells. A comparative analysis between MCF7 611-CTF and MCF7 p185 cells revealed increased chromatin accessibility at genes involved in some key oncogenic pathways, such as the Akt pathway (*AKT1* and *HSP90AA1*) and MAPK pathway (*MAPKAPK2*, *HRAS*, and *CACNB3*, among others), as well as several PAM50 genes (*CDH3*, *BIRC5*, and *CCNE1*). *TP73*, a regulator of p53, is also highly open in MCF7 611- CTF cells^44^, which could be potentially consistent with the higher aggressivity of the tumors expressing this proteoform. Conversely, several genes showed reduced accessibility in MCF7 611- CTF cells compared to the p185. These include *NFIB*, a gene responsible for the remodeling of chromatin in small cell lung cancer^45^; *KLF4*, a tumor suppressor^46^; *TRIM14*, whose downregulation inhibits cell proliferation while enhancing apoptosis^47^; or *FRK*, a kinase associated with cell proliferation, migration, and invasiveness^48^.

### EpiBlot reveals extensive chromatin remodeling and protein expression changes in chemotherapy-treated HER2 proteoform-expressing cells

Drug treatments can profoundly alter cell fate and are known drivers of tumor evolution^49^. We decided to test EpiBlot on doxorubicin-treated cell lines. Doxorubicin is a broadly-used anthracycline that functions by sequestering DNA topoisomerase II to create DNA double-strand breaking points^50,51^, leading to apoptosis of the cells (**Supplementary** Figure 8A). In addition to inducing DNA damage, doxorubicin evicts histones from specific genomic regions^52^, and alters nucleosomal occupancy levels on cancer related genes, potentially promoting genomic instability^13^. Given the impact that doxorubicin has on chromatin, we expected to observe widespread changes before and after treatment across the different cell lines.

We first induced HER2 expression in the HER2-proteoform transfected cell lines through the incorporation of doxocycline to the culture medium, and then we treated the cells with 1 μM of doxorubicin for 48 h. To determine the effects of doxorubicin on chromatin accessibility, we performed plateATAC-seq on treated and untreated samples from each cell line. To assess whether doxorubicin-induced changes were biologically stochastic, we performed plateATAC-seq on three biological replicates per condition of the treated cells and performed PCA (**Supplementary** Figure 9A). Both in treated and untreated conditions, the three replicates cluster together, indicating similarity in the chromatin remodeling profiles. Treated and untreated conditions were clearly separated for all the three conditions, with MCF7 611-CTF presenting the greatest separation. In contrast, MCF7 parental and MCF7 p185 were clustered more closely together. An example of the changes in chromatin accessibility is shown in **Figure 2A**, where the *ERBB2* gene shows decreased accessibility following treatment across all conditions. The impact of doxorubicin treatment on chromatin accessibility on selected genes is visualized in **Figure 2B**. *CDK6* exhibits increased accessibility only in the doxorubicin-treated condition, while *PIK3R2* is selectively closed in MCF7 611-CTF doxorubicin-treated samples but remains accessible in other cell lines and conditions. Differential accessibility analysis reveals that doxorubicin treatment has an extensive impact in chromatin accessibility (**Figure 2C)**. MCF7 611-CTF exhibits the highest number of differentially accessible regions (∼64,768), followed by MCF7 p185 (∼48,785) and MCF7 parental (∼34,017), suggesting a stronger chromatin remodeling response in MCF7 611- CTF. Differentially accessible regions (FDR-adjusted p-value < 0.001) with significant chromatin accessibility changes include *CCND1*, which is downregulated (more closed chromatin) in doxorubicin-treated MCF7 611-CTF, suggesting suppression of cell-cycle regulation. *GATA3* shows closed accessibility in doxorubicin-treated MCF7 p185, potentially indicating downregulated transcription factor activity. *AZIN1-AS1* exhibits greater accessibility in doxorubicin-treated MCF7 parental and MCF7 p185 cells, while remaining unchanged in MCF7 611-CTF (**Figure 2C**). The gene ontology enrichment pathway analysis shows that MCF7 611- CTF has a strong enrichment in pathways related to epithelial-mesenchymal transition, *ERBB2* signaling, and cell adhesion, whereas doxocyclin-treated MCF7 p185 is enriched in pathways related to RNA splicing, histone modification, and *Wnt* signaling (**Figure 2D**). Shared pathways include *ERBB2* signaling, *MAPK* cascade regulation, and DNA-binding transcription factor activity, suggesting common and distinct chromatin accessibility responses to doxorubicin in these HER2 proteoforms.

**Figure 2.**
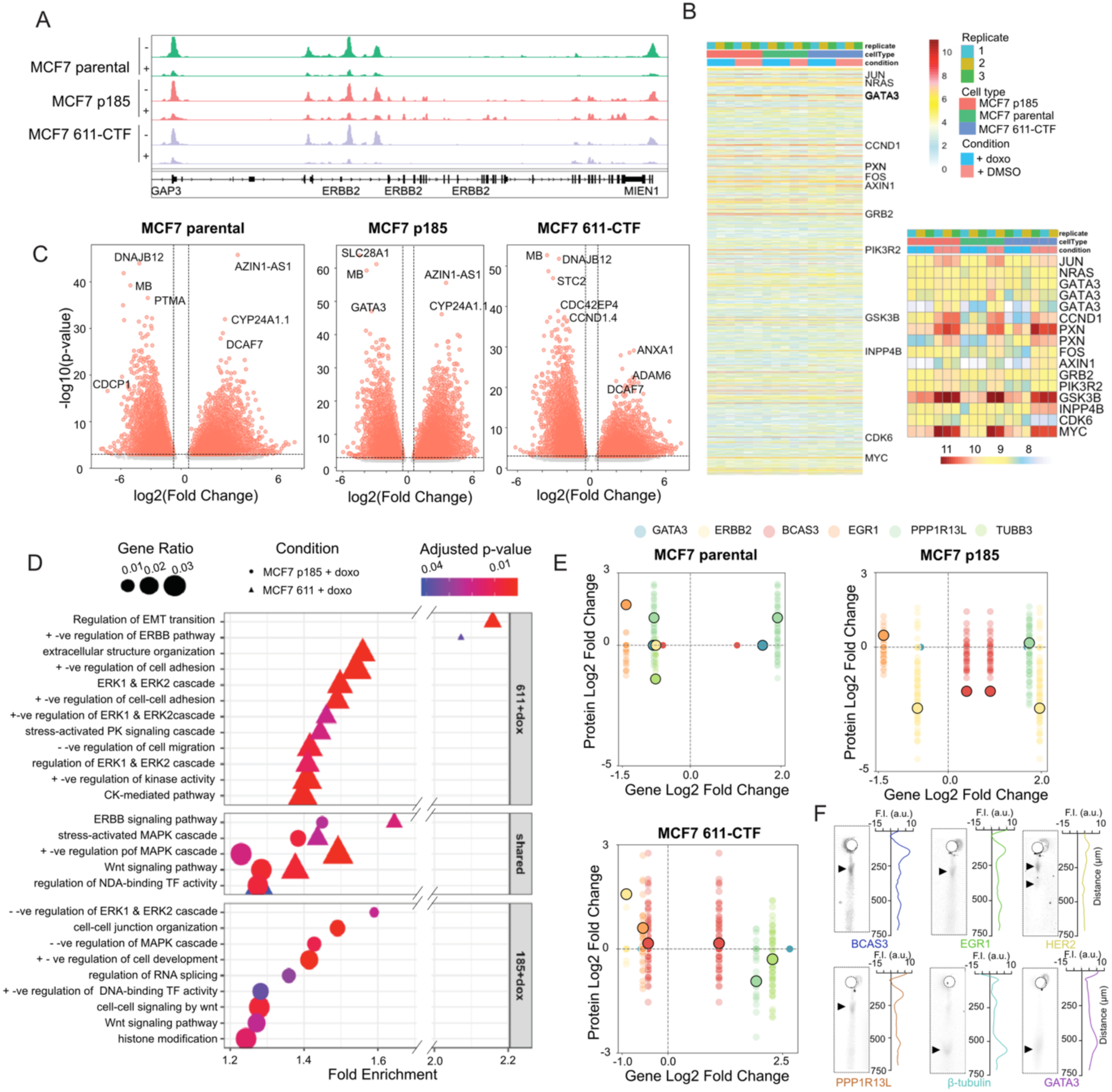
Doxorubicin treatment induces widespread changes in chromatin accessibility across all cell types. A) Changes in chromatin accessibility at the ERBB2 gene in MCF7 parental, p185, and 611-CTF cell lines with (+) and without (-) doxo treatment. Genome browser tracks (GRCh38) show chromatin accessibility at the ERBB2 (HER2) and MIEN1 gene loci in MCF7-transfected cell lines with and without doxo. Differentially accessible (DA) peaks are suppressed in the ERBB2 and MIEN1 region in all doxo-treated samples compared to control samples within the cell lines. All track lines have the same y-axis limits, and peak heights are scaled across all samples. B) Heatmap of differential chromatin accessibility in key breast cancer-related genes across MCF7-derived cell lines (parent, p185, and 611-CTF) treated with doxo or control. Data were Z-score normalized per gene (row normalization) to highlight relative accessibility changes. Doxo treatment results in widespread chromatin closure, with MYC, PXN, AXIN1, JUN, GATA3, and CCND1 showing reduced accessibility across all cell lines. Samples are clustered based on chromatin accessibility, with colors representing Z-scores (red = increased accessibility, blue = decreased accessibility). C) Volcano plots showing differentially accessible chromatin regions between doxo-treated and control conditions in parent, p185, and 611-CTF MCF7-derived cell lines. Each point represents an individual chromatin region, with red indicating significantly differentially accessible regions (FDR-adjusted p-value < 0.001) and grey representing nonsignificant regions. The x-axis (Log₂ fold change) indicates accessibility changes, where negative values (left) represent regions more accessible in control, and positive values (right) represent regions more accessible in doxo-treated samples. The vertical dotted line represents an absolute log2 fold change of 0.5, while the horizontal dotted line represents an adjusted p-value threshold of 0.001. FDR-corrected p-values were obtained using DESeq2. D) Gene ontology (GO) enrichment analysis of differentially accessible chromatin regions in doxorubicin-treated MCF7 p185 (185+doxo) and MCF7 611-CTF (611+doxo) cell lines. Enriched pathways are categorized into those specific to 611+doxo (top panel), shared between MCF7 611+doxo and MCF7 185+doxo (middle panel), and specific to 185+doxo (bottom panel). The x-axis represents fold enrichment, while the y-axis lists significantly enriched GO terms. Dot size corresponds to the gene ratio, and color indicates the adjusted p-value, with red representing the most significant pathways (lowest p-value). For brevity on y-axis labels, ‘positive’ is abbreviated as “+-ve”, and negative is abbreviated as “ - -ve”. E) Scatterplot of Protein Log2 fold change and gene Log2 fold change after doxo treatment. Circle with a dark outline represents the average Log2 fold change across all the tested single cells. F) Single-cell Western blot of selected proteins in MCF7 611-CTFcells after doxo treatment. False-color micrographs show the protein location, with companion intensity trace at the right side of each. F.I. corresponds to fluorescence intensity and a.u. to arbitrary units.

Following the plateATAC-seq analysis, we tested by scWB a selection of protein markers chosen based on their relevance to cancer biology and changes in the chromatin accessibility profiles (**Supplementary** Figure 10). Apart from its archiving capability, the use of scWB offers several additional advantages: sample material requirements are notably less than a conventional WB, and single-cell resolution reports heterogeneity of protein expression among a cell population. We chose the targets GATA3, a transcription factor implicated in the luminal differentiation of breast epithelium^53^; EGR1, a protein with both tumor-suppressor and tumor-promoter activities^54^; BCAS3, associated with chemoresistance through a p53-mediated mechanism^55^; and PPP1R13L, implicated in tumorigenesis^56^ (**Figure 2E-F, Supplementary** Figure 10**&11A**). Additionally, we included HER2 and β-tubulin as controls. PCA of single-cell protein expression profiles shows that treated and untreated cells form partially overlapping clusters, indicating that the protein-level differences are not completely distinct (**Supplementary** Figure 10B). The partial separation is a product of multiple proteins showing significantly different expression between the treated and non-treated expression (*P*<0.05, ANOVA test), especially in the case of MCF7 p185 (**Supplementary** Figure 10C). Despite changes in both chromatin accessibility and protein abundance, we observed a wide dynamic range of protein expression across individual cells, a hallmark of a heterogeneous response to doxorubicin (**Figure 2E**).

### Targeted treatment induces limited changes

While doxorubicin produced extensive changes in the cell profile, we anticipated that a targeted treatment would result in more moderate changes. For that, we tested the effects of the tyrosine kinase inhibitor lapatinib. Lapatinib inhibits the phosphorylation of the intracellular tyrosine kinase domain of HER2 (**Supplementary** Figure 8B). Given its structure, the drug remains effective against full and truncated forms of HER2 such as 611-CTF^14,57^. Nonetheless, HER2 611-CTF forms homodimers maintained by disulfide bonds^58^, potentially influencing downstream signaling and response to kinase inhibition.

We induced the cells via doxocycline and followed with lapatinib treatment. The treatment had a modest impact on global chromatin accessibility. The number of differentially accessible regions was limited: 23 regions for MCF7 611-CTF, followed by MCF7 parental with 87 regions, and MCF7 p185, with 159. PCA revealed that PC1, which accounts for 82% of the total variance, separates samples primarily by cell line, whereas PC2 (8% variance) captures differences between treatment conditions (**Supplementary** Figure 9B). The dominant contribution of PC1 indicates that HER2 proteoform status exerts a stronger influence on chromatin accessibility than lapatinib treatment under these conditions. For instance, the *ERBB2* locus showed minimal accessibility changes following treatment across all three cell lines (**Figure 3A**), and no strong differences were observed based on treatment (**Figure 3B**). The low number of differentially accessible regions suggests a limited effect of lapatinib on global chromatin remodeling. Nonetheless, a direct comparison of the treated cell lines MCF7 p185 and 611-CTF reveals widespread changes in chromatin accessibility (**Figure 3C**). This finding highlights how the expression of HER2 proteoforms can profoundly alter the cellular response to therapy. Through a gene ontology enrichment analysis, we observe MCF7 611-CTF-specific pathways include *ERBB* signaling, *Wnt* signaling, and ERK1/2 cascade regulation, suggesting a strong enrichment of HER2-related pathways in MCF7 611-CTF under lapatinib treatment (**Figure 3D-E**). In contrast, MCF7 p185- specific pathways involve MAPK activity and Ras protein signaling, as previously reported in HER2+ cells^59,60^, and TOR signaling, a pathway involved in lapatinib resistance^61,62^. These differences in downstream signaling responses to lapatinib suggest distinct effects of the two HER2 proteoforms. Shared pathways include MAPK cascade regulation and Wnt signaling, highlighting common chromatin accessibility changes between the two cell lines.

**Figure 3.**
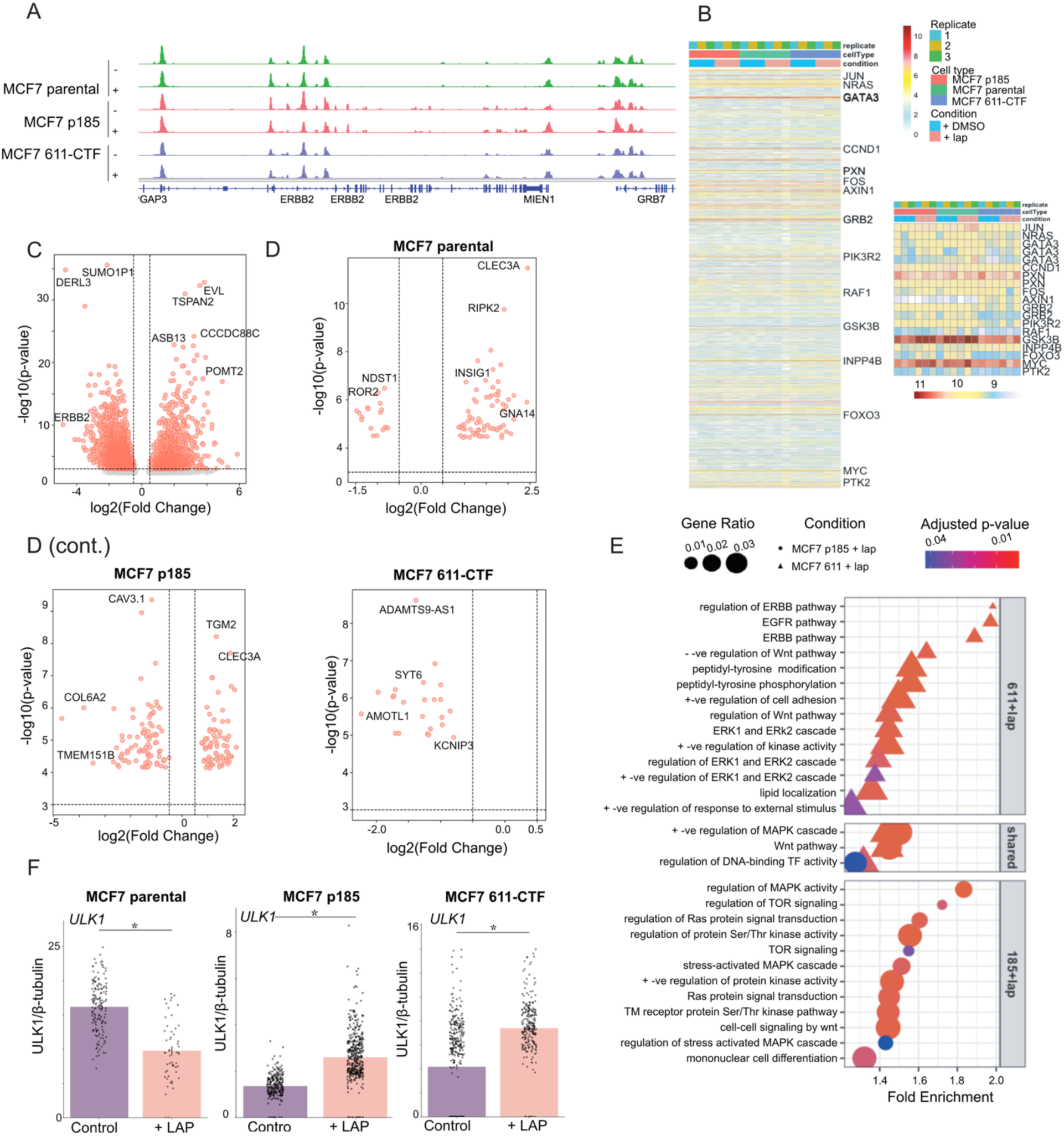
Lapatinib treatment leads to modest changes in chromatin. A) Changes in chromatin accessibility at the ERBB2 gene in MCF7 parental, p185, and 611-CTF cell lines with (+) and without (-) lapatinib treatment. Genome browser tracks (GRCh38) show chromatin accessibility at the ERBB2 (HER2), MIEN1, and GRB7 gene loci in MCF7-transfected cell lines with and without lapatinib treatment. Lapatinib show minimal to no effect between the rearment in each cell line. All track lines have the same y-axis limits, and peak heights are scaled across all samples. B) Heatmap of chromatin accessibility in 100 key breast cancer-related genes across MCF7- derived cell lines (parental, p185, and 611-CTF) treated with lapatinib (lap) or control (DMSO). Data were Z-score normalized per gene (row normalization) to highlight relative accessibility changes. A subset heatmap zooms in on regions with at least one condition exceeding a Z-score of 9.5, with further scaling applied to enhance visualization. C) Volcano plot of differentially accessible chromatin regions between MCF7 611-CTF and MCF7 p185 cell lines under lapatinib (lap) treatment. Each point represents a chromatin region, with red indicating significant changes (FDR-adjusted p-value < 0.00001). The x-axis (Log₂ fold change) shows regions more accessible in p185 (left) or 611-CTF (right), while the y-axis (-log₁₀ p-value) represents significance. D) Volcano plots of differentially accessible chromatin regions between lapatinib-treated (lap) and control (DMSO) conditions in parental, p185, and 611-CTF MCF7 cell lines. Each point represents a chromatin region, with red indicating significant changes (FDR-adjusted p-value < 0.0001). The x-axis (Log₂ fold change) shows regions more accessible in control (left) or lap-treated (right), while the y-axis (-log₁₀ p-value) represents significance. Lapatinib induces minimal chromatin accessibility changes across all cell lines, with p185 showing the strongest response and 611-CTF exhibiting the weakest. The vertical dotted line represents an absolute log2 fold change of 1.0, while the horizontal dotted line represents an adjusted p-value threshold of 0.00001. FDR- corrected p-values were obtained using DESeq2. E) GO enrichment analysis of differentially accessible chromatin regions in 611-CTF (611lap) and p185 (185lap) cell lines under lapatinib treatment. Enriched pathways are categorized into those specific to 611lap (top panel), shared between both cell lines (middle panel), and specific to 185lap (bottom panel). The x-axis represents fold enrichment, while the y-axis lists significantly enriched GO terms. Dot size corresponds to the gene ratio, and color indicates the adjusted p-value, with red representing the most significant pathways. For brevity on y-axis labels, ‘positive’ is abbreviated as “+-ve”, and negative is abbreviated as “ - -ve”. F) Bar plot showing ULK1 expression increase after lapatinib treatment in MCF7 p185 and 611- CTF, while presenting a significant decrease in MCF7 parental cells.

Based on the results of the plateATAC-seq analyses, we scrutinized protein targets (**Figure 3F, Supplementary** Figure 12C) in individual cells by scWB. We selected ULK1, a protein that induces autophagy^63^, and TFAP2C, a regulator of estrogen receptor expression in breast cancer^64^. Only two proteins were selected given the limited number of changes observed. As control proteins, we once again included HER2 and β-tubulin. MCF7 parental shows a decrease in expression of ULK1, which corresponds well to the decrease in open regions for the *ULK1* gene (**Figure 3F, Supplementary** Figure 11B**&12**). Although no changes were observed in the differential accessibility in MCF7 p185 and 611-CTF, we did observe a significant increase in ULK1 expression in both cell lines. Similarly, an increase was observed in TFAP2C abundance in MCF7 p185, although the TFAP2 locus closed after treatment.

### Identification of subpopulations and divergent accessibility patterns in tumor organoids

To test the compatibility of EpiBlot with clinical samples, we used patient-derived organoids. Organoids are frequently used tumor models, capable of preserving the cellular heterogeneity and overall complexity of the tumors from which they are derived^35^, making them more physiologically relevant systems. In this study, we examined a cohort of 2 breast cancer PDOs and one normal breast PDO, derived from tumor-adjacent non-tumorous normal breast tissue for comparison (**Figure 4A**). We specifically selected TNBC organoids to investigate the behavior of HER2-low phenotypes in a relevant context, given the emerging significance of HER2 expression gradients in TNBC. The TNBC organoids were confirmed to be HER2 low by immunofluorescence (**Supplementary** Figure 13).

**Figure 4.**
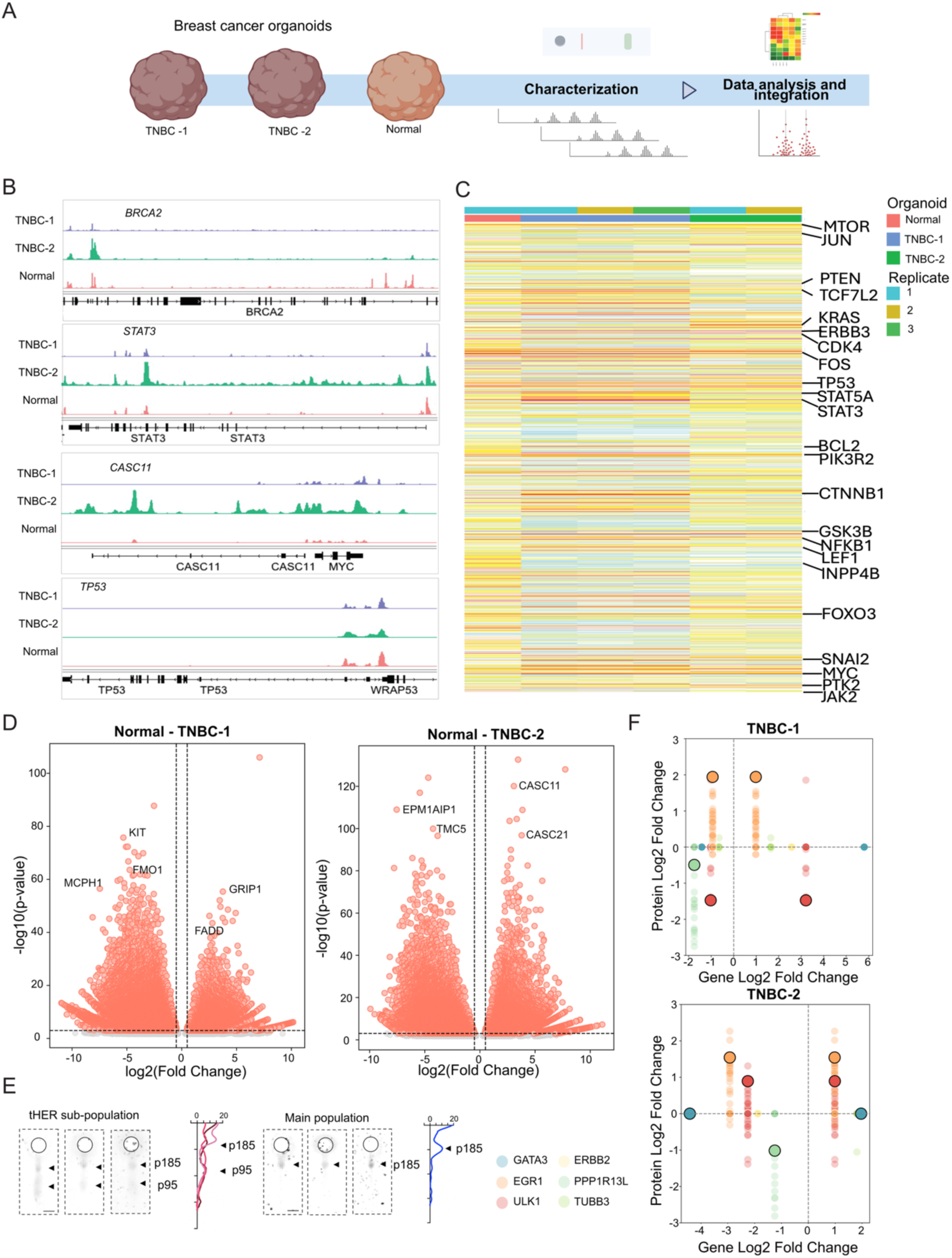
Chromatin accessibility and HER2 protein expression in TNBC patient-derived organoids. A) Schematic representation of triple-negative breast cancer (TNBC) organoids derived from Grade 2 (TNBC-2) and Grade 3 (TNBC-1) human tumors, alongside a normal human breast tissue organoid (Normal). The workflow includes protein characterization, chromatin accessibility profiling, and integrative data analysis. B) Genome browser tracks showing chromatin accessibility profiles at key breast cancer- associated loci (BRCA2, STAT3, CASC11, MYC, TP53, and WRAP53) across TNBC-1, TNBC- 2, and Normal. C) Heatmap of chromatin accessibility across key signaling genes in TNBC organoids and normal controls. Each row represents a genomic region, with color intensity reflecting accessibility levels (Z-score normalized). Warmer colors (red/yellow) indicate higher accessibility, while cooler colors (blue/green) represent lower accessibility. Annotated regions with Z-score > 9.5 are labeled, highlighting regions with extreme chromatin accessibility changes. D) Volcano plots of differentially accessible chromatin regions in TNBC-2 (left) and TNBC-1 (right), using Normal as the reference. Each point represents a chromatin region, with red indicating significant changes (FDR-adjusted p-value < 0.001). E) Micrograph of the scWB bands expressing the truncated p95HER2 proteoform (n=3, left), compared to three bands expressing the main proteoform (n=869, right). Undetermined stains are shown in Supplementary Figure 15. F) Protein log2Fold change compared to gene log2Fold change of TNBC-1 and TNBC-2 using Normal as reference. Circle with a dark outline represents the average log2fold across all the tested single cells.

Dissociated organoids were divided in two fractions, one submitted for plateATAC-seq and the other reserved for scWB. Technical replicates of TNBC-1 and TNBC-2 clustered together (**Supplementary** Figure 9C), and all organoids were distinct from one another. The selected loci in **Figure 4B** show that TNBC-1 exhibits strong chromatin accessibility upstream of *BRCA2* and *STAT3*, while TNBC-2 has the most open chromatin in the upstream region. At *CASC11*, TNBC- 2 displays the highest accessibility, whereas Normal and TNBC-1 shows a closed pattern. In the *TP53-WRAP53* region, TNBC-1 exhibits the strongest chromatin accessibility, potentially reflecting an altered regulatory state in Grade 3 compared to Grade 2 (TNBC-2) and normal tissue. Compared to Normal PDOs (due to the limited number of cells only one replicate was analyzed), TNBC organoids have fewer accessible promoter regions (≤1kb) and more distal intergenic regions, suggesting a shift toward noncoding regulatory elements in TNBC (**Figure Supplementary 7B**). **Figure 4C** shows that TNBC-2 exhibits an opposite accessibility pattern compared to both normal controls and TNBC-1, suggesting a distinct chromatin remodeling profile in this TNBC subtype. Notably, genes involved in cancer-related pathways, including *MTOR*, *JUN*, *STAT3*, *MYC*, *TP53*, and *CTNNB1*, show differential chromatin accessibility, indicating potential regulatory divergence in TNBC-2 compared to other TNBC models. Both cancer organoids present a downregulation of *ERBB2* compared to the Normal organoid, despite showing an enrichment the MAPK pathway (gene ratio 0.04) in TNBC-1, and PI3K-Akt and MAPK in TNBC-2 (gene ratio 0.05 and 0.045) (**Supplementary** Figure 14). Both TNBC-1 and TNBC-2 exhibit distinct chromatin accessibility profiles compared to Normal (**Figure 4D**). TNBC-1 shows more closed chromatin regions relative to Normal, suggesting widespread chromatin compaction, while TNBC-2 exhibits a more balanced mix of open and closed regions, indicating differential chromatin remodeling between TNBC organoid models.

We evaluated the protein expression of GATA3, ULK1, EGR1, PPP1R13L, HER2, and β-tubulin across the three sequenced organoids, based on the previous selection of markers (**Supplementary 15A**). We did not observe a distinctive clustering of either of the organoids based on protein markers (**Supplementary** Figure 15B). Interestingly, we detected a small cell subpopulation (n=3, i.e., 0.3% of examined cells) in TNBC-1, which expressed the truncated HER2 proteoform (**Figure 4E**). The number of truncated cells could be higher, since we observed several scWB bands with a weak signal in the molecular weight compatible with truncated HER2 (n=26, **Supplementary** Figure 15C). However, we discounted these weak bands from the analysis as the signal was below the quantification limit. The reduced size of the observed subpopulation could be caused by inadvertent bias in sampling into the two aliquots required for the two different analyses, the low levels of expression of the truncated HER2 proteoforms, or the reduced size of the subpopulation. Additionally, we observed that the expression of HER2 in both TNBC organoids was significantly higher (*P*<0.05) than in the Normal tissue, despite presenting a decreased accessibility in the locus, possibly caused by changes in the accessibility of the enhancer region of *ERBB2* (**Supplementary** Figure 14C and 15D).

Finally, we examined the correlation between protein and chromatin accessibility by comparing the Log2 fold-change for each marker (**Figure 4F**). In both tumorous organoids, EGR1 expression is elevated, while two main peaks exist for the chromatin accessibility, one increasing 1-fold and the other decreasing 3-fold. ULK1 shows an opposite tendency for TNBC-1 and TNBC-2, with TNBC-2 increasing the expression of this protein.

## Discussion

EpiBlot is a new pooled-cell assay, which combines plateATAC-seq for chromatin accessibility analysis and a single-cell resolution protein target assay, scWB. PlateATAC-seq profiles chromatin accessibility with limited numbers of starting nuclei, while scWB archives proteins for a subsequent targeted analysis on the same experimental sample, mitigating batch effects. Our findings demonstrate that HER2 proteoforms exert a potent influence on cellular regulation. As an upstream receptor tyrosine kinase, HER2 plays a pivotal role in signal transduction, and variations in its expression or activity have the potential to determine cell fate. Naïve MCF7 cells are generally classified as HER2 0 or +1 ref.^65^. Overexpression of full-length HER2 resulted in relatively modest alterations to the chromatin accessibility landscape, with only 7 differentially accessible genes. In contrast, expression of a truncated, constitutively active HER2 proteoform produced a substantial shift in chromatin architecture, yielding between 100 and 2,400 unique differentially accessible regions when compared to MCF7 parental and p185. This underscores the profound regulatory impact of specific proteoform identity beyond overall HER2 expression levels.

One of the key advantages of EpiBlot is the archiving of proteins from the same experiment for a later examination once sequencing results are available. This approach minimizes the need for additional experiments to test proteins of interest. Protein expression reflects downstream biological states, and their expression is generally expected to correlate with upstream transcriptional and epigenetic regulation. However, previous studies have shown only moderate correlations between protein levels, RNA abundance, and chromatin accessibility^16,22^. Our results align with this pattern, with the observed variance exacerbated by single-cell heterogeneity. Differential chromatin accessibility did not always translate into corresponding protein changes. This may be due to additional layers of epigenetic regulation, lack of functional relevance of the epigenetic changes, or differences in protein synthesis and degradation dynamics^66^. Future studies may consider temporally resolved measurements of chromatin accessibility and protein levels to delineate the dynamics of proteomic responses following chromatin remodeling, especially at the single cell level.

Cell-line identity emerged as a key determinant in response to chemotherapeutic treatment. Doxorubicin, a DNA intercalating agent, induced widespread changes in chromatin accessibility across all cell lines. These effects are likely attributable to the drug’s known mechanism of drug action, which involves chromatin intercalation and disruption of DNA topology. The potential impact of HER2 amplification on anthracycline efficacy has been debated, with some studies reporting enhanced therapeutic responses in HER2+ tumors^67–70^ and highlighting sensitization of p95HER2/611-CTF-expressing xenografts to trastuzumab^12^. However, a recent meta-analysis has failed to establish a consistent link between anthracycline administration and complete clinical response in HER2+ patients^71^.

As anticipated for a targeted therapy, lapatinib had a more limited effect on chromatin accessibility. Interestingly, MCF7 611-CTF cells exhibited fewer differentially accessible genes. The clinical significance of p95HER2 expression in lapatinib-treated patients remains unclear, with some reports associating it with longer survival^72^ and others finding no significant impact^73^. These discrepancies likely reflect differences in study design, treatment regimens, and patient populations.

In our analysis of the PDOs, we observed a predominance of closed chromatin regions in tumor samples relative to the normal control (open vs closed: TNBC-1 35k vs 47k, and TNBC-2 32k vs 44k). This is consistent with findings in other disease states where global chromatin compaction was observed, such as macular degeneration^74^ and systemic sclerosis^75^. These findings may suggest that the repression of regulatory elements may be a feature of TNBC epigenomes. Notably, despite the overall decrease in chromatin accessibility at the *ERBB2* locus, both TNBC organoids exhibited elevated HER2 protein levels, suggesting that post-transcriptional regulation or enhancer-driven expression may contribute to HER2 upregulation in these tumors. These findings highlight that, while it may appear that there is epigenetic silencing of the *ERBB2* gene, alternative regulatory mechanisms are inducing HER2 expression. This protein upregulation may facilitate the uptake of HER2-directed drugs into the cancer cells.

Intratumor heterogeneity can be captured in PDOs^36^, and we could capture such heterogeneity in TNBC-1, which presents a small subpopulation of cells expressing p95HER2. Tumor heterogeneity in HER2+ tumors has been widely reported^76–79^, and its presence in HER2 proteoform expression highlights the importance of multi-modal single-cell analysis capable of simultaneously assessing proteoform expression and chromatin accessibility, as the detected subpopulation of cells may be expressing a distinct chromatin accessibility pattern. The detection of HER2 proteoforms in a TNBC-classified organoid suggests the need of use of personalized treatments adequate to the development of tumor heterogeneity of each patient. The relevance of understanding intra-tumor heterogeneity in TNBCs has been widely explored, showing correlation to metastasis^80^. The ability to detect such rare subclones could be pivotal for stratifying patients for therapy or identifying resistance mechanisms. However, higher-throughput, integrated single-cell technologies that can simultaneously profile chromatin and proteoforms will be essential to fully capture tumor complexity. For clinical applications, such systems would benefit of the examination of nuclear characteristics as samples derived from patients may contain fibroblasts and immune cells among tumor cells.

A limitation of the technological aspect in plateATAC-seq is that, while the study is performed on a scant number of cells (300-500 cells), the initial concentration of nuclei required is high to allow the visualization of the cell pellet. Similarly, scWB presents constraints in protein analysis, with target multiplexing requiring multiple rounds of stripping and probing to analyze >3 protein targets, a process that can alter epitope availability^34^. Furthermore, while the single-cell resolution of a scWB is an advantage to understand tumor heterogeneity, reliable detection is restricted to the top half of the mammalian proteome^30^.

Together, our study reveals that HER2 proteoform diversity alters chromatin organization and modulates drug response, with truncated forms exerting more profound effects than full-length HER2.

## Acknowledgements

Photolithography was performed in the QB3 Biomolecular Nanotechnology Center at UC Berkeley, imaging was performed at the Cell & Tissue Analysis Facility at UC Berkeley, and the sequencing at the Genomics Platform at the San Francisco CZ Biohub. We thank Dr. Alison Su and Dr. Angelika Feldmann for helpful discussions. We thank Joaquin Arribas, PhD and his lab for providing the MCF7 transfected cells.

## Supplementary data

Supplementary data is available.

## Conflict of interest

None declared.

## Funding

A.E.H. and J.M.R. acknowledge financial support from the Investigator Program of the Chan Zuckerberg Biohub San Francisco. A.E.H. acknowledges support from the US National Institutes of Health (NIH R01CA20301). J.M.R. thanks the Dana-Farber Cancer Institute Breast Biospecimen Users Committee, DF/HCC Breast SPORE 1P50CA168504, NIH 5R01CA281361, and METAvivor. A.F.K. acknowledges support from the Swiss National Science Foundation (SNSF) Postdoc Mobility fellowship (P500PN_214236). The content is solely the responsibility of the authors and does not necessarily represent the official views of the National Institutes of Health/NCI.

## Notes

### Competing Interest Statement

The authors have declared no competing interest.

### Summary of Updates

Manuscript was adjusted to clarify the impact of the presented method.

