## Supplementary Information for "EpiBlot: Joint Mapping of Chromatin Accessibility and Targeted Proteomics in HER2-expressing Breast Cancer Systems"

**Supplementary Figure 1: Discontinuation of ab16901 during this study and substitution by ab134182.**

The authors of this study are aware of the reproducibility crisis driven by unspecific antibodies and aim to provide a transparent use and justification of antibody choice.

The antibody ab16901 was initially chosen due to its compatibility with HER2 proteoforms. During the course of this study, abcam discontinued the previously used ab16901, as in a recent analysis (end of 2024) with HER2 knock-off cell lines, this antibody showed nonspecific signal using immunohistochemistry on formalin-fixed paraffin-embedded cells. abcam offered a substitution antibody (ab134182), which was formulated against the same antigen as ab16901.

We proceeded to test this new antibody using the scWB. As the reader might be aware, WBs are a different analytical tool commonly used to validate antibodies. The results can be seen in the figure.

While we repeated our experiments on transfected MCF7 cell lines with the new antibody, we finally decided to use the data obtained with the discontinued antibody:

1) The MCF7 cell lines that we use were provided by the Arribas lab, who already characterize such cell lines[7]. When performing the scWB experiments, we observe bands at the expected molecular weights for p185 and CTF-611. We do not observe unspecific bands using ab16901, while we do see them in ab134182.
2) No discovery studies have been performed with the discontinued antibody. Organoids were tested with the new antibody formulation. While the final intensity of detection is less in the new antibody, the PCA plots are performed with Z-normalization, reducing the impact of signal intensity.

The image below shows the comparison between the two antibodies. The top image shows the same slide, first stained with ab16901, then stripped and restained with ab134182. The bottom image shows two different slides, one stained with each antibody (first staining).

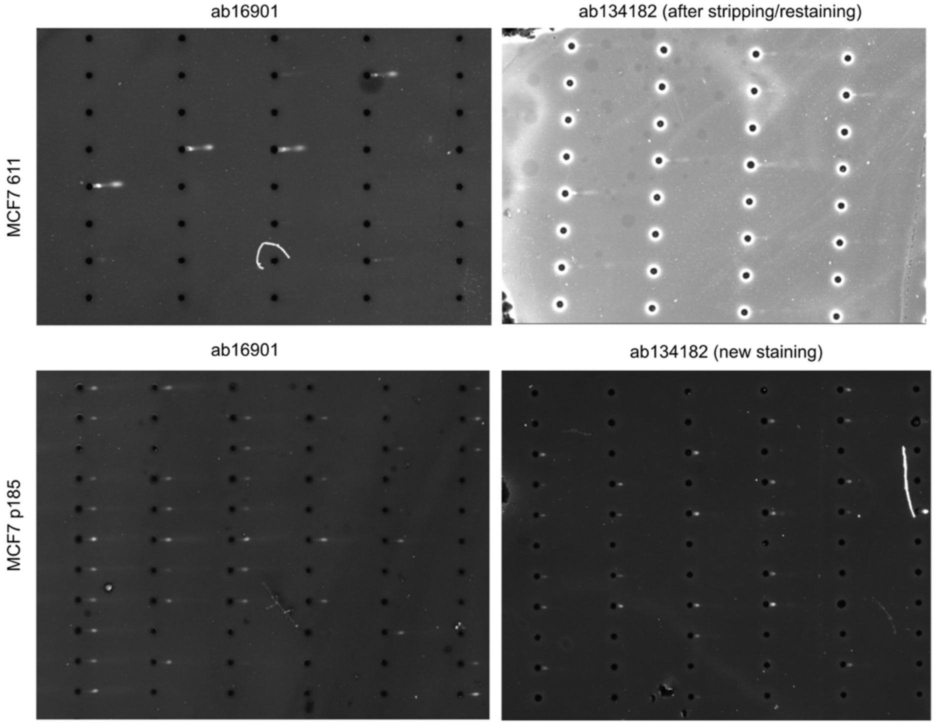

**Supplementary Figure 2: Data analysis pipeline for protein segmentation.**
CNN: convolutional neural network.

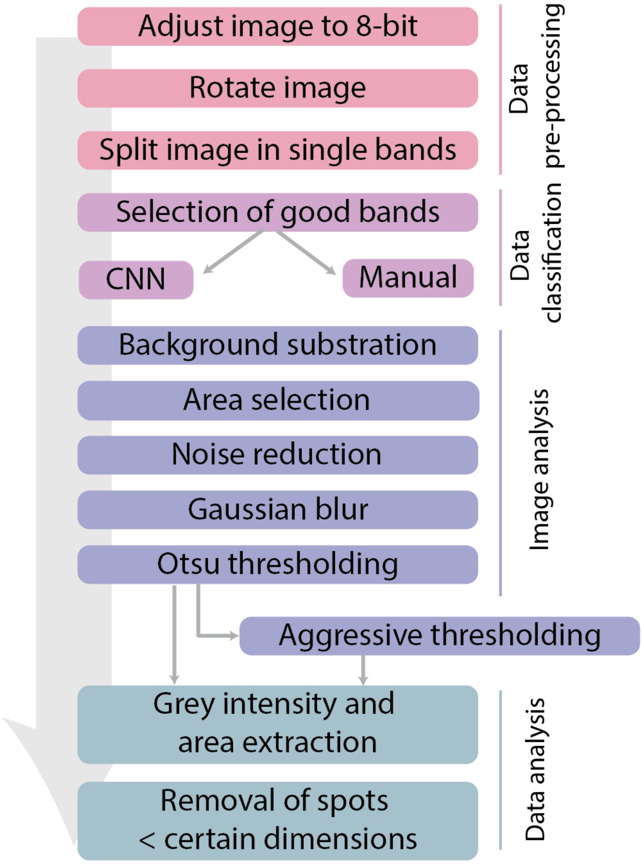

**Supplementary Figure 3: Fluorescence images stained for HER2, F-actin, and nuclei in transfected MCF7 cell lines.**

A) MCF7 parental (empty vector), B) MCF7 p185, and C) MCF7 611-CTF. Note that the controls for doxorubicin and lapatinib differ in DMSO concentration, as each control was prepared with the respective volume of DMSO. Scale bar: 20 µm.

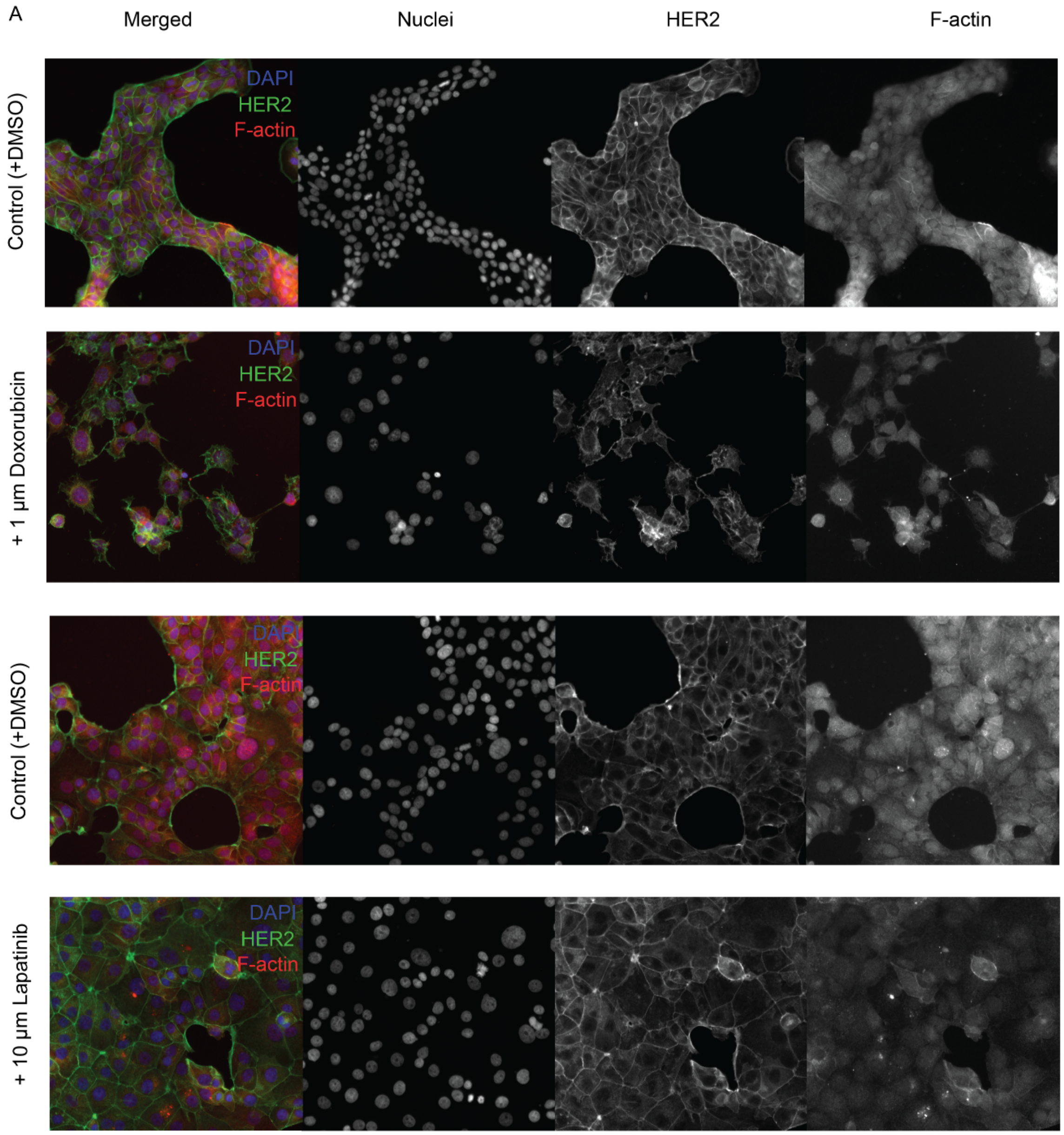

**Supplementary Figure 3** (cont.)

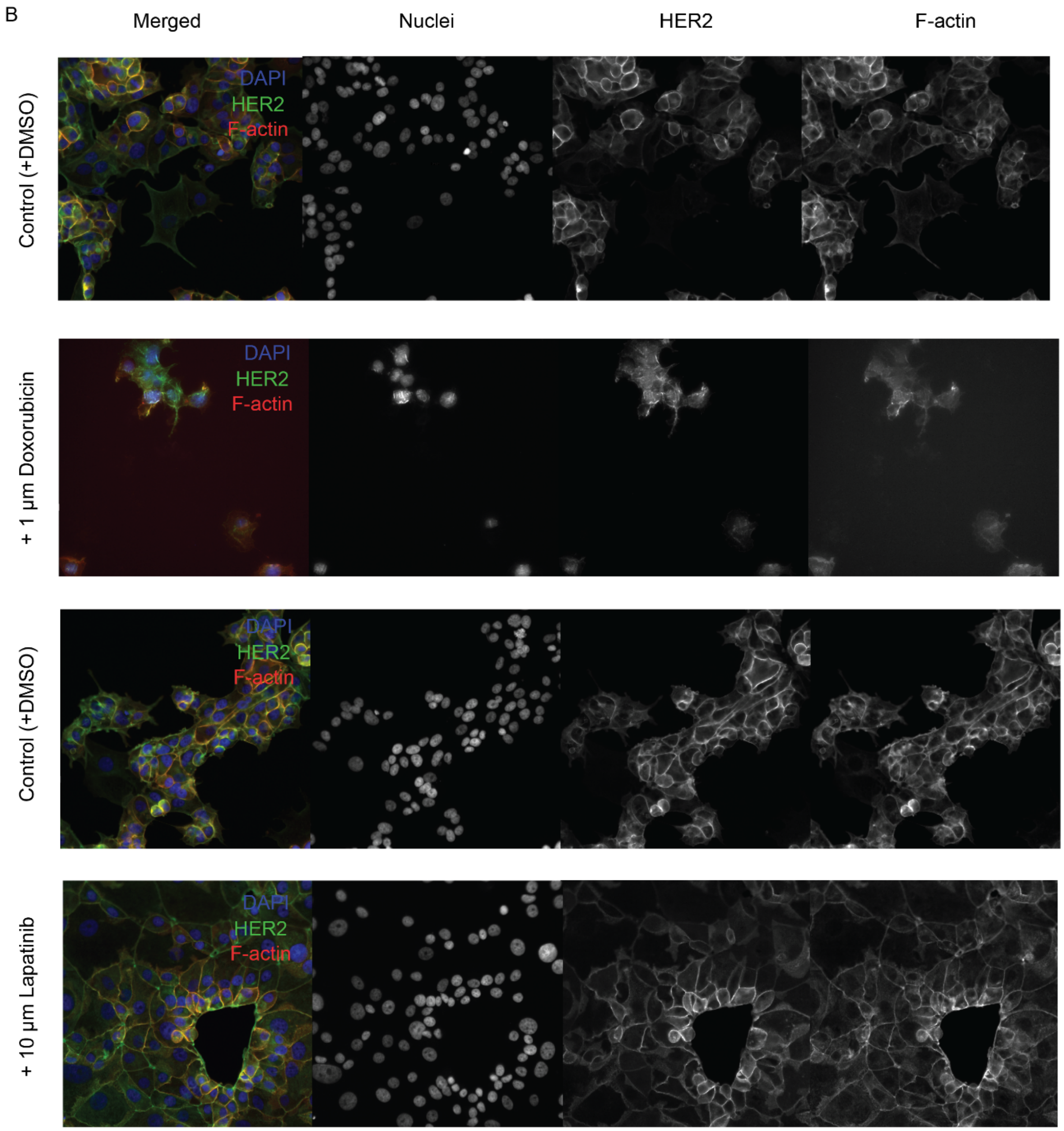

**Supplementary Figure 3** (cont.)

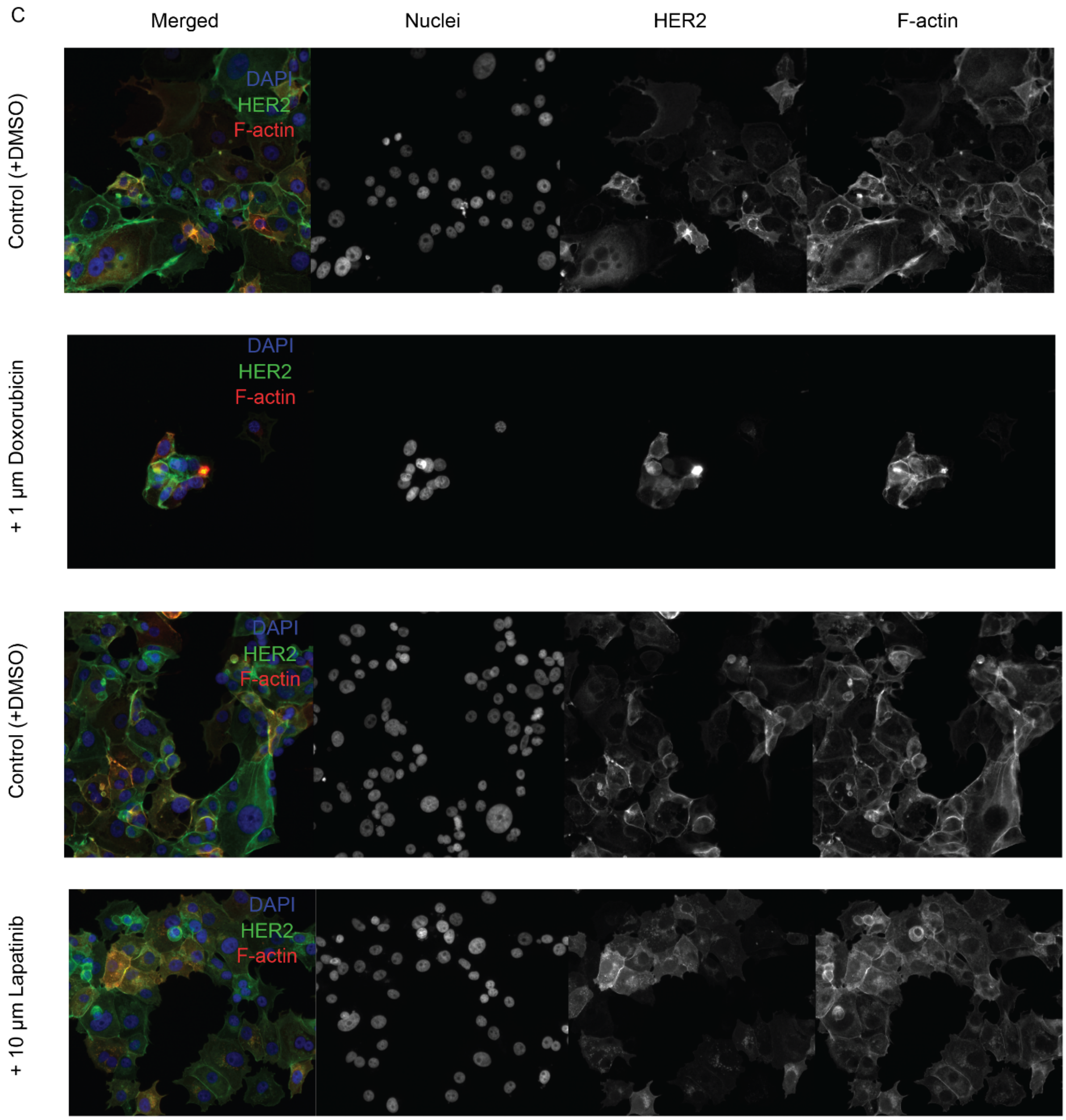

**Supplementary Figure 4: Analysis of nuclear dimensions.**
Area, circularity, length, mean intensity inside of the nucleus and ratio between the area of the nucleus and the area of the cytoplasm for A) the three cell lines (MCF7 parental, MCF7 p185, and MCF7 611-CTF), B) the same cell lines before and after doxorubicin treatment, and C) before and after lapatinib treatment. Significance: **** is $p < 0.0001$, *** is $p < 0.001$, ** is $p < 0.01$, and * is $p < 0.05$. Non-significance is not plotted. Statistics in B and C were calculated only pairwise between no treatment (-) and treatment (+).

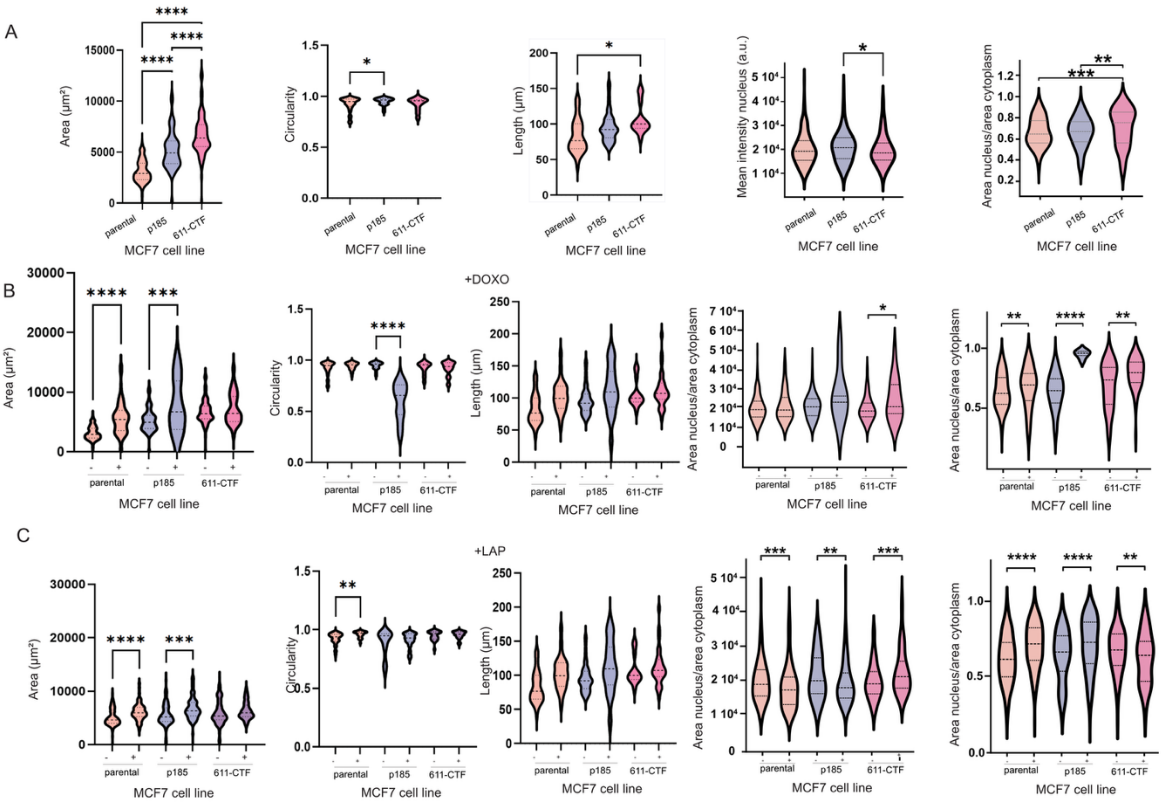

**Supplementary Note 1: Analysis of nuclear dimensions.**

As part of the characterization of the cell lines, we performed an analysis of the nuclear morphology. Nuclear morphology can provide information about biological states such as aneuploidy, genomic instability, transcriptional dysregulation, and genetic mutations[1]. Further, nuclear morphology in cancer, including differences in nuclei that have been observed between different subtypes of breast cancer, has been shown to correlate with diagnosis[1–3]. To explore morphological changes associated with HER2 overexpression, we stained the nuclei with Hoechst 33342 to quantify nuclear area, circularity and roundness. Given that the sole variable across our conditions was HER2 proteoform expression, we aimed to determine whether the expression of these proteoforms influences nuclear architecture.

*Nuclear changes after doxorubicin treatment*

The nuclear area increased significantly ($p<0.0001$, MCF7 parental $n=233$, MCF7 p185 $n=115$, and MCF7 611-CTF $n=104$ nuclei) with HER2 overexpression, and this effect was further amplified in cells expressing the 611-CTF fragment. When normalized by the cytoplasm area, we observe no changes between MCF7 parental and MCF7 p185, but there is an increase in nuclear area in MCF7 611-CTF, as confirmed by the increased nuclear length ($p<0.5$). Although we observed significant differences ($p<0.5$) in circularity, length, and grey intensity of some of the nuclei, no trends were observed. To the best of our knowledge, the mechanistic basis of nuclear enlargement remains unclear. In breast cancer, nuclear size, atypical features, and pleomorphism are included in standard grading systems, with higher nuclear grades correlating with worse overall prognosis[5]. In this case, we observe the largest nuclear areas in MCF7 611-CTF cells, which correspond to aggressive tumors[7].

Both MCF7 parental and MCF7 p185 presented a significant nuclei enlargement following doxorubicin treatment (MCF7 parental $n_{DMSO}=233$, $n_{DOXO}=75$; MCF7 p185 $n_{DMSO}=115$, $n_{DOXO}=17$; MCF7 611-CTF $n_{DMSO}=104$, $n_{DOXO}=28$, $P \leq 0.0001$, $P \leq 0.001$, and not significant, respectively), while MCF7 611-CTF remained similar. If we observe the ratio between nuclear and cytoplasmic area, all of the conditions showed an increase after treatment, indicating that the nucleus increased in area more than the cytoplasm.

*Nuclear changes after lapatinib treatment*

The morphological analysis of nuclei showed a significant increase in nuclear area in MCF7 parental and MCF7 p185 following lapatinib treatment (MCF7 parental $n_{DMSO}=194$, $n_{LAP}=156$; MCF7 p185 $n_{DMSO}=90$, $n_{LAP}=225$, $P \leq 0.0001$, $P \leq 0.001$, respectively), despite the inhibition of the pathway. This tendency is maintained if the ratio between nucleus and cytoplasm is taken into consideration. This was not the case in MCF7 611-CTF (MCF7 611-CTF $n_{DMSO}=151$, $n_{LAP}=101$, not significant), although when we evaluated the ratio, we observed a decrease in area after treatment. Interestingly, we observed that DAPI intensity was significantly increased in MCF7 611-CTF, despite no consistent size changes on the nucleus, while opposite was true for the other two cell lines.

We identified significant changes in nuclear morphology, specifically nuclear area, associated with both HER2 proteoform expression and drug treatment. While nuclear area is influenced by multiple factors, including chromatin compaction, mechanical tension, and osmotic pressure, evidence argues against a direct correlation with chromatin condensation alone[4]. Following doxorubicin treatment, we observed a downregulation of LMNA and ACTB across all cell lines, which may contribute to increased nuclear drug uptake and a consequent increase in nuclear size[6]. A similar, though less pronounced, enlargement was observed after lapatinib treatment. Although specific studies on nuclear size changes induced by lapatinib are limited, it is plausible that shared mechanisms involving cytoskeletal or nuclear envelope alterations may contribute to a nuclear enlargement.

**Supplementary Figure 5. Optimization of the plateATAC-seq protocol.**

A) Schematic representation of the plateATAC-seq workflow. A total of 300 nuclei were treated with a 1:15 Tn5 mix in a 15 µL microwell plate at 37°C for 30 minutes, followed by Tn5 release using EDTA to quench MgCl2 at 55°C for an additional 30 minutes. Library preparation via indexing PCR was subsequently performed. B) Characterization of EDTA for releasing Tn5 and determination of optimal PCR cycles for library preparation. C) Illustration of the final ATAC-seq pooled library, incorporating data from 58 libraries derived from four transfected MCF7 cell lines (parental, 185, and 611), each under different treated conditions including control, DMSO as control for doxorubicin-treated condition, doxorubicin, DMSO as control for lapatinib-treated condition, and lapatinib.

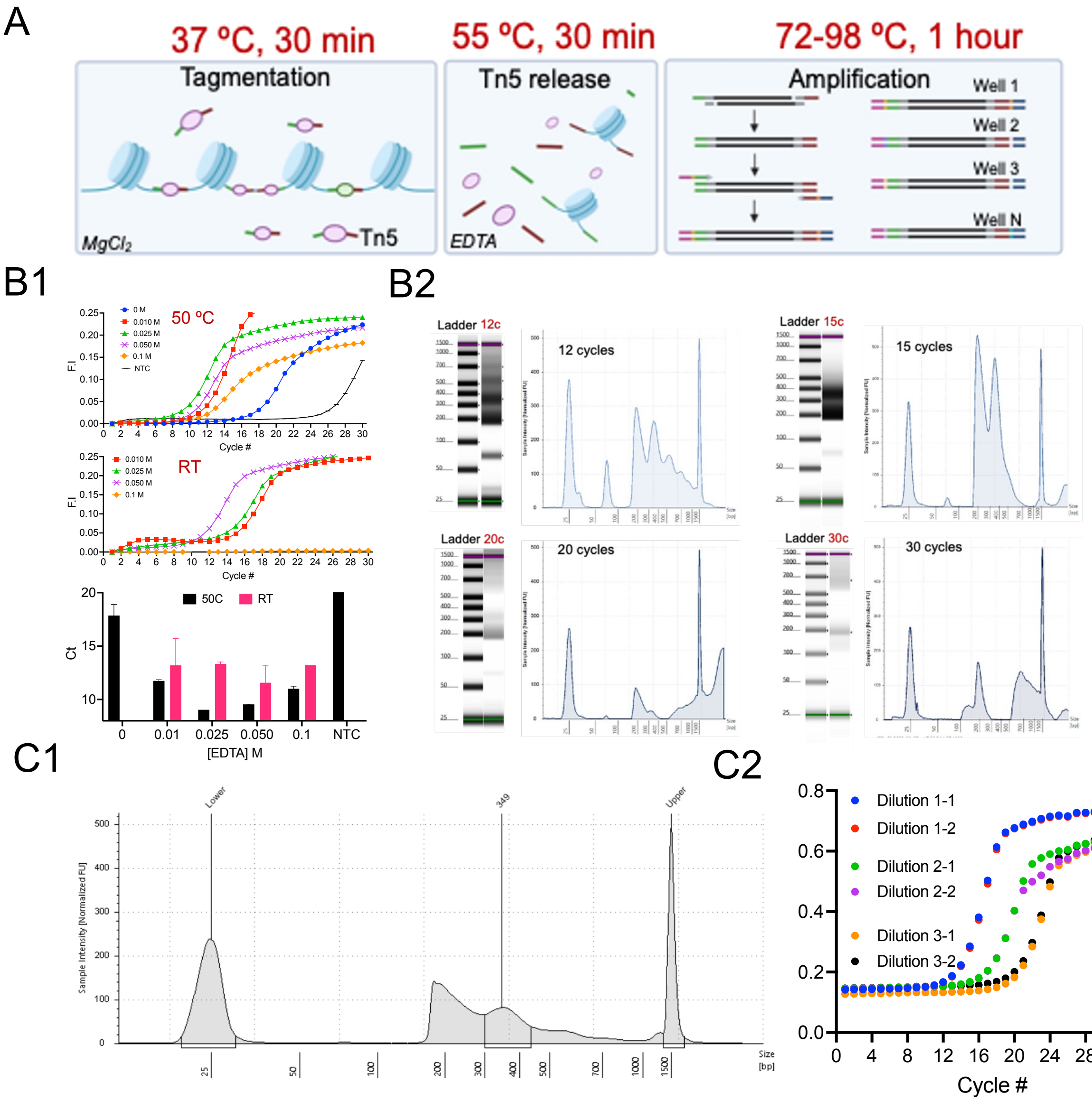

**Supplementary Figure 6. Sequencing metrics QC of the plate-based ATAC method.**
A) Comparison of library size estimated by the Picard tool, fraction of mitochondrial DNA (MT content), and fraction of reads in peaks (FRiP) in plate-based ATAC-seq libraries from MCF7 transfected cells. The plate-based method yielded nearly 60% FRiP and <20% mitochondrial content across samples. B) Comparison of peak profiles in bulk ATAC-seq with MCF7 parental (50,000 nuclei) and plate-based ATAC (300 nuclei) in chr16 (378 kb) (Top) and in the ERBB2 locus at chr17 q12 (50kb) (bottom). C) Insert size reads across 48 samples of MCF7 transfected cell lines with different treatments (no DMSO, DMSO as control for doxorubicin-treated condition, doxorubicin, DMSO as control for lapatinib-treated condition, and lapatinib). D) ATAC-seq metaplot illustrating the genomic distribution of open chromatin regions relative to transcription start sites (TSS). The x-axis delineates the distance from the TSS, ranging from -3000 to 3000 base pairs, with 0 representing the TSS. The y-axis indicates the occurrence or density of ATAC-seq reads. A dashed red line denotes the position of the TSS.

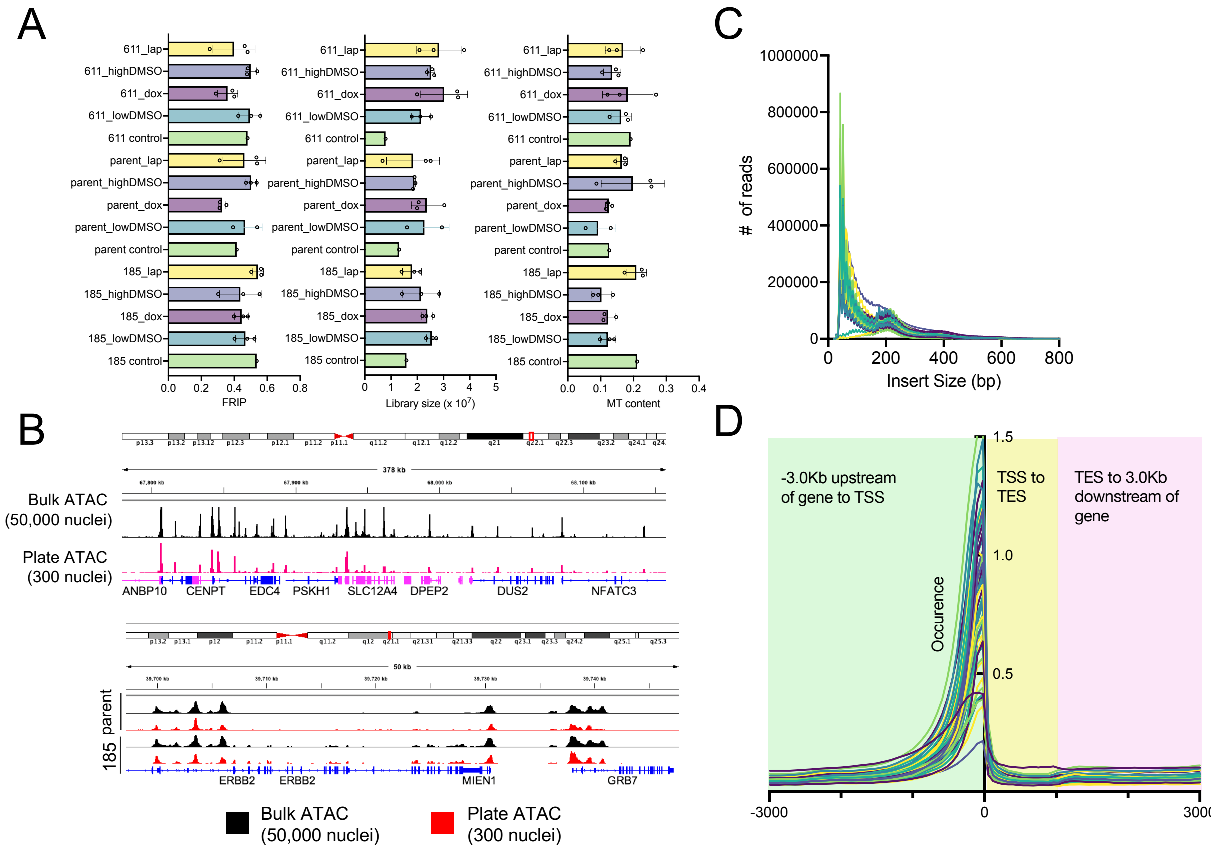

**Supplementary Figure 7: Genomic distribution of differentially accessible (DA) chromatin regions.**

A) Genomic distribution of DA regions for MCF7 transfected cell lines. Pie charts showing the percentage of DA ATAC-seq peaks located in promoter, intronic, intergenic, and exonic regions for the three MCF7-transfected cell lines expressing different HER2 proteoforms. B) Genomic distribution of differentially accessible chromatin regions in Normal, TNBC-1 and TNBC-2 organoids. Pie charts represent the percentage of differentially accessible regions mapped to various genomic features, including promoters, untranslated regions (UTRs), exons, introns, downstream regions, and distal intergenic regions.

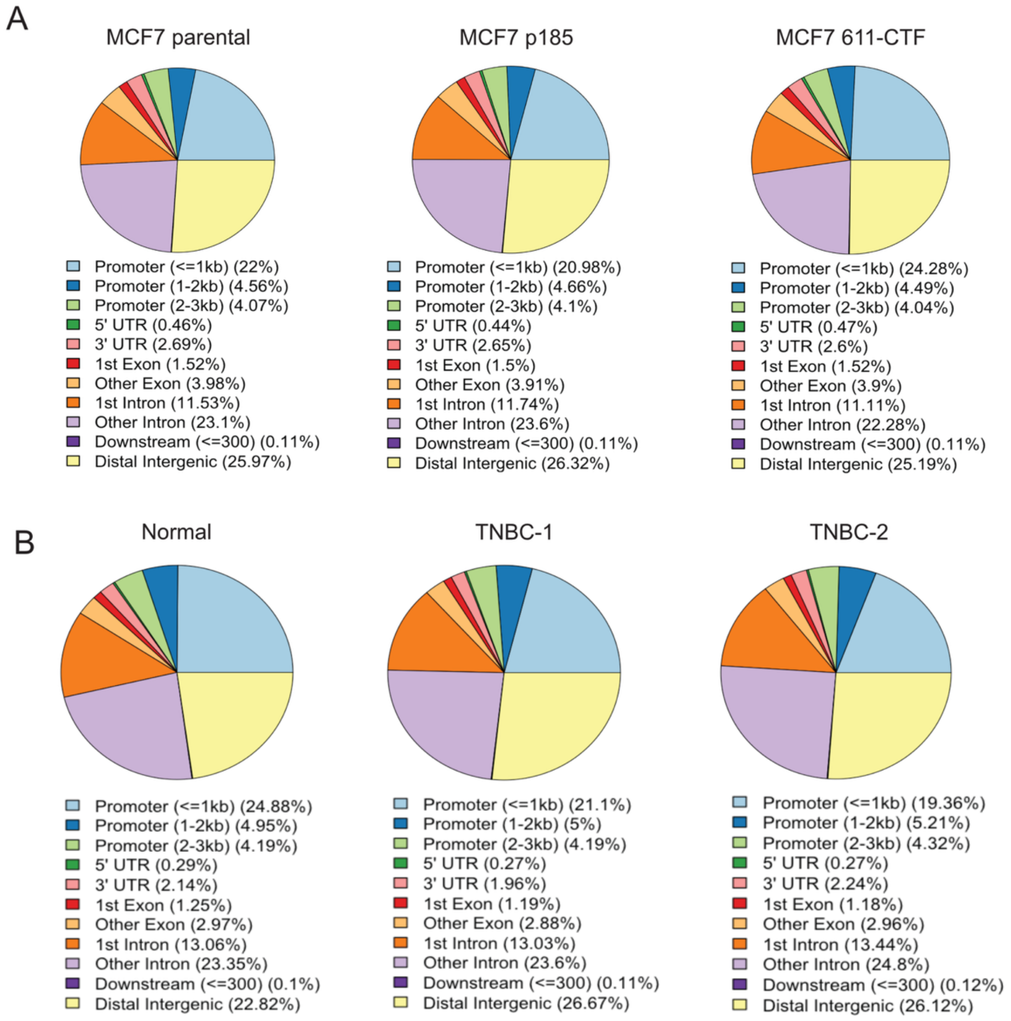

**Supplementary Figure 8. Mechanism of action of selected drugs.**

A) Scheme showing the mechanism of action of doxorubicin (DOXO) on the chromatin of the cells1. DDR: DNA damage response. B) Scheme representing the mechanism of action of lapatinib (LAP). Lapatinib inhibits the tyrosine kinase domain of HER2, inhibiting PI3K/Akt and Ras/MAPK pathways2.

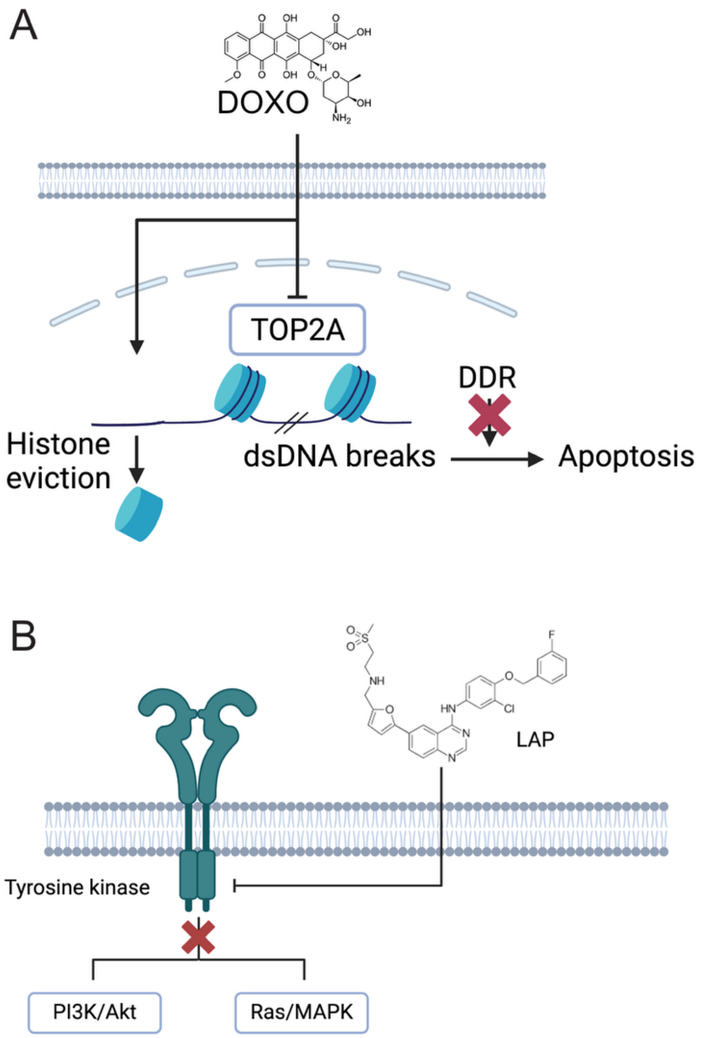

**Supplementary Figure 9. PCA of the different conditions.**

A) PCA of ATAC-seq signal in MCF7-derived cell lines treated with doxorubicin (doxo) or control (DMSO). PC1 (82% variance) primarily separates doxo-treated from control samples, indicating that doxo treatment is the major driver of chromatin accessibility changes. PC2 (8% variance) captures additional variation, likely reflecting differences between cell lines. p185+doxo (red) and parental+doxo (green) cluster closely together, suggesting a similar chromatin accessibility response to doxorubicin. In contrast, 611-CTF+doxo (blue) clusters separately, indicating a distinct chromatin remodeling response compared to p185 and parental under doxorubicin treatment. B) PCA of chromatin accessibility in MCF7 cell lines (parental, p185, and 611-CTF) treated with lapatinib (lap) or control (DMSO). PC1 (75% variance) separates 611-CTF from p185 and parent, indicating distinct chromatin accessibility profiles between these HER2 proteoforms. PC2 (7% variance) captures minimal variation due to treatment, suggesting that lapatinib induces only subtle chromatin accessibility changes in p185 and parent cells. MCF7 611-CTF shows no clear separation between lap-treated and control conditions, indicating a lack of response at the chromatin level. Overall, cell line identity is the primary driver of variation, while lapatinib treatment has a minimal effect on chromatin accessibility. C) PCA of chromatin accessibility in the tested organoids. PC1 (90% variance) primarily separates the organoids, indicating major chromatin accessibility differences. TNBC-1 (blue) is the most distinct, clustering farthest from the others, suggesting a highly altered chromatin landscape. TNBC-2 (green) is the most similar within its group, forming a tight cluster on the right. Normal (red) is intermediate, positioned separately along PC2 (10% variance), suggesting greater variability compared to TNBC-2 but less distinct than TNBC-1. The strong PC1 separation highlights major epigenetic differences among these TNBC organoid models.

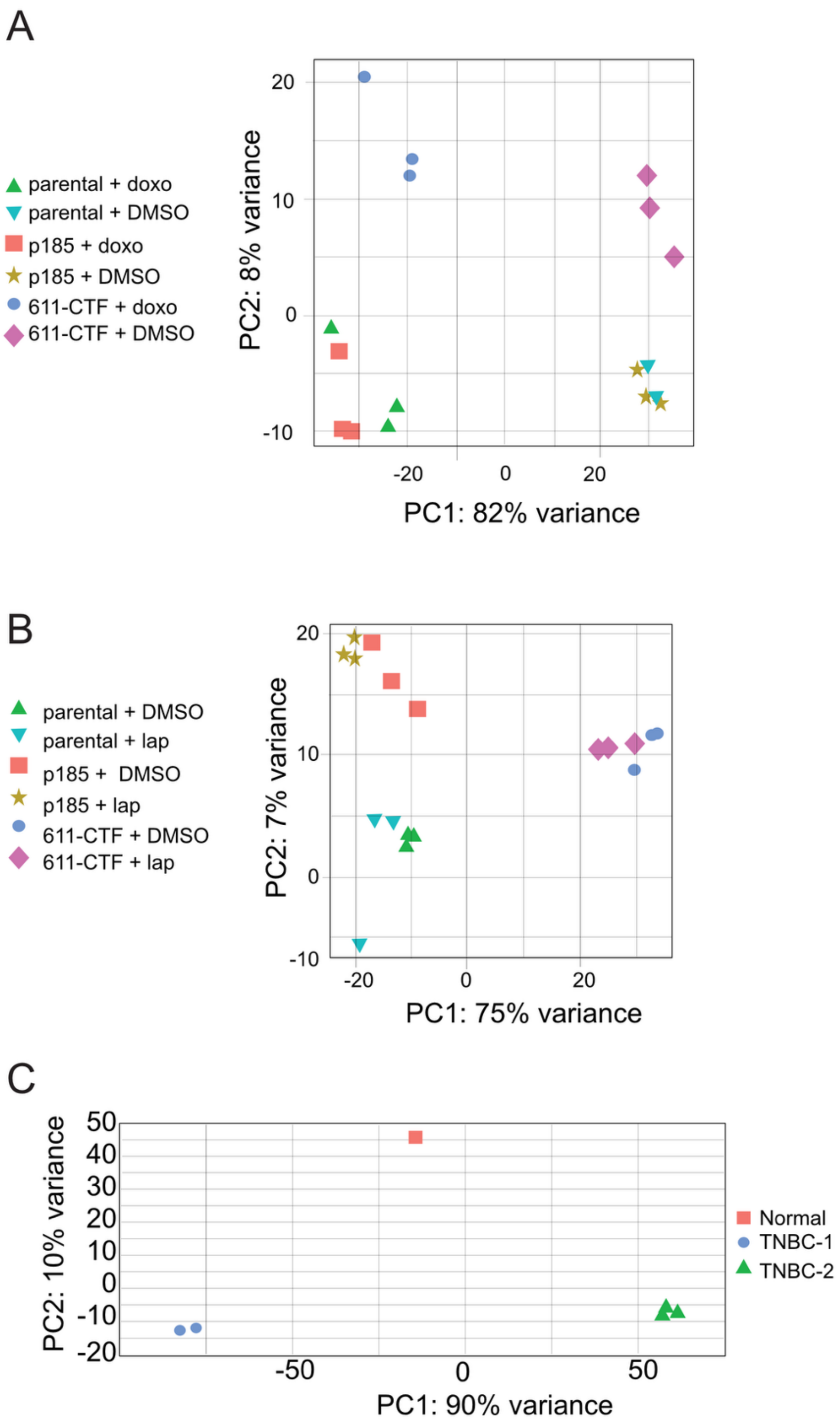
A
parental + doxo
parental + DMSO
p185 + doxo
p185 + DMSO
611-CTF + doxo
611-CTF + DMSO
20
10
0
-10
PC2: 8% variance
-20
0
20
PC1: 82% variance
B
parental + DMSO
parental + lap
p185 + DMSO
p185 + lap
611-CTF + DMSO
611-CTF + lap
20
10
0
-10
PC2: 7% variance
-20
0
20
PC1: 75% variance
C
50
40
30
20
10
0
-10
-20
PC2: 10% variance
-50
0
50
PC1: 90% variance
Normal
TNBC-1
TNBC-2

**Supplementary Figure 10: Protein analysis in transfected MCF7 cell lines upon doxorubicin treatment.**

A) Location of the tested protein on the volcano plot (DMSO control vs doxorubicin treated). B) PCA plot of the proteins divided into treated and untreated conditions. C) Bar blot with the selected tested proteins (ERBB2, EGR1, PPP1R13L, GATA3, BCAS3) on the 3 cell lines. Conditions with no expression were not plotted. The y-axis represents the protein expression level normalized by the expression of β-tubulin in the same cell. Proteins labeled with [Form 1 or 2] presented proteoforms in these gels. Form 1 corresponds to the observed proteoform with higher molecular weight, while Form 2 is the lower molecular weight form. Significant differences were shown as * (P<0.05).

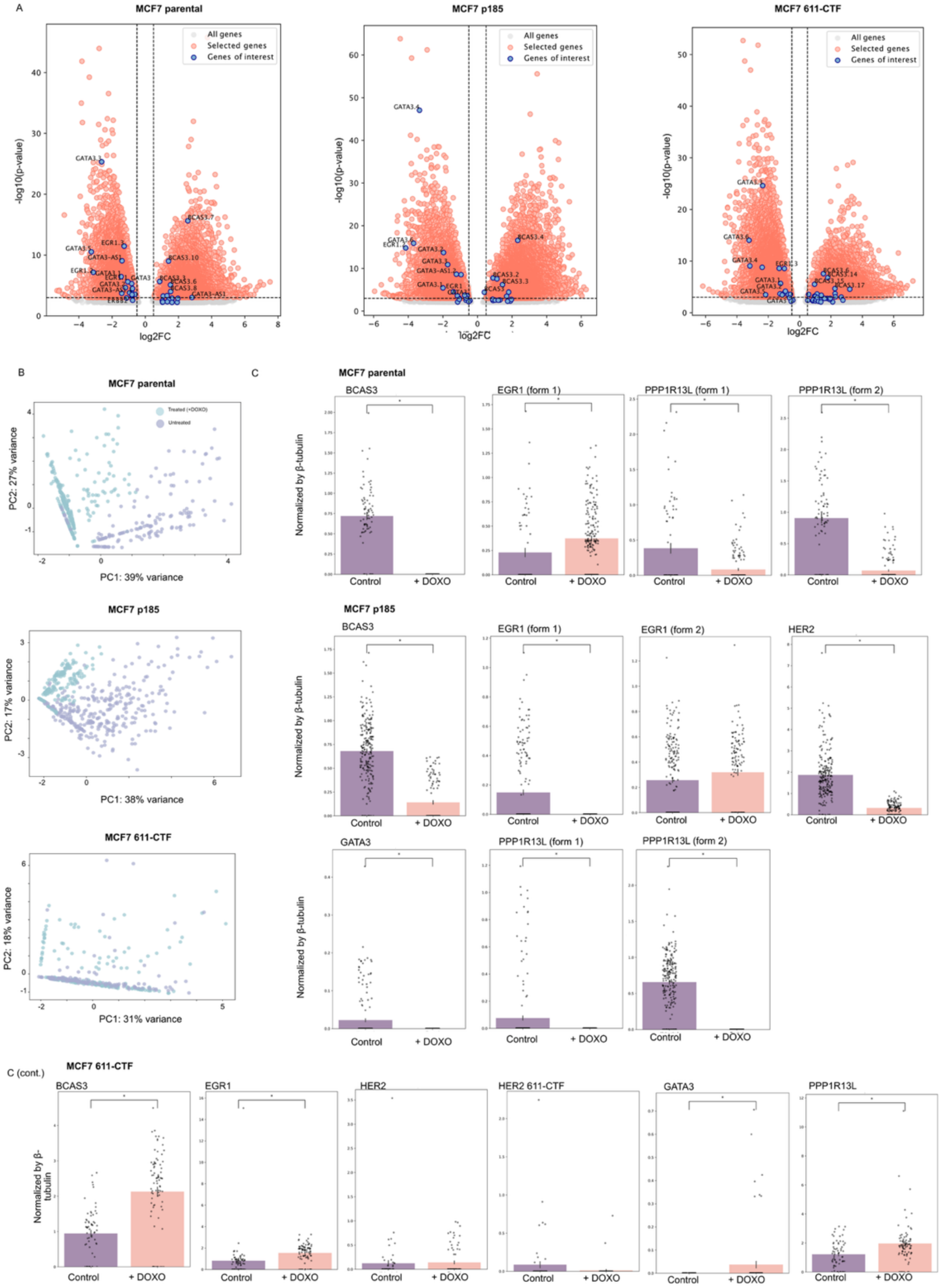

**Supplementary Figure 11: Changes in chromatin accessibility between several conditions.** A) Comparison of MCF7 parental, p185, and 611-CTF before and after doxorubicin treatment for GATA3, BCAS3, EGR1, and PPP1R13L. B) Comparison of MCF7 parental, p185, and 611-CTF before and after lapatinib treatment for TFAP2C and ULK1. C) Comparison between the Normal, TNBC-1 and TNBC-2 organoids for proteins ERBB2, PPP1R13L, ULK1, EGR1, and GATA3.

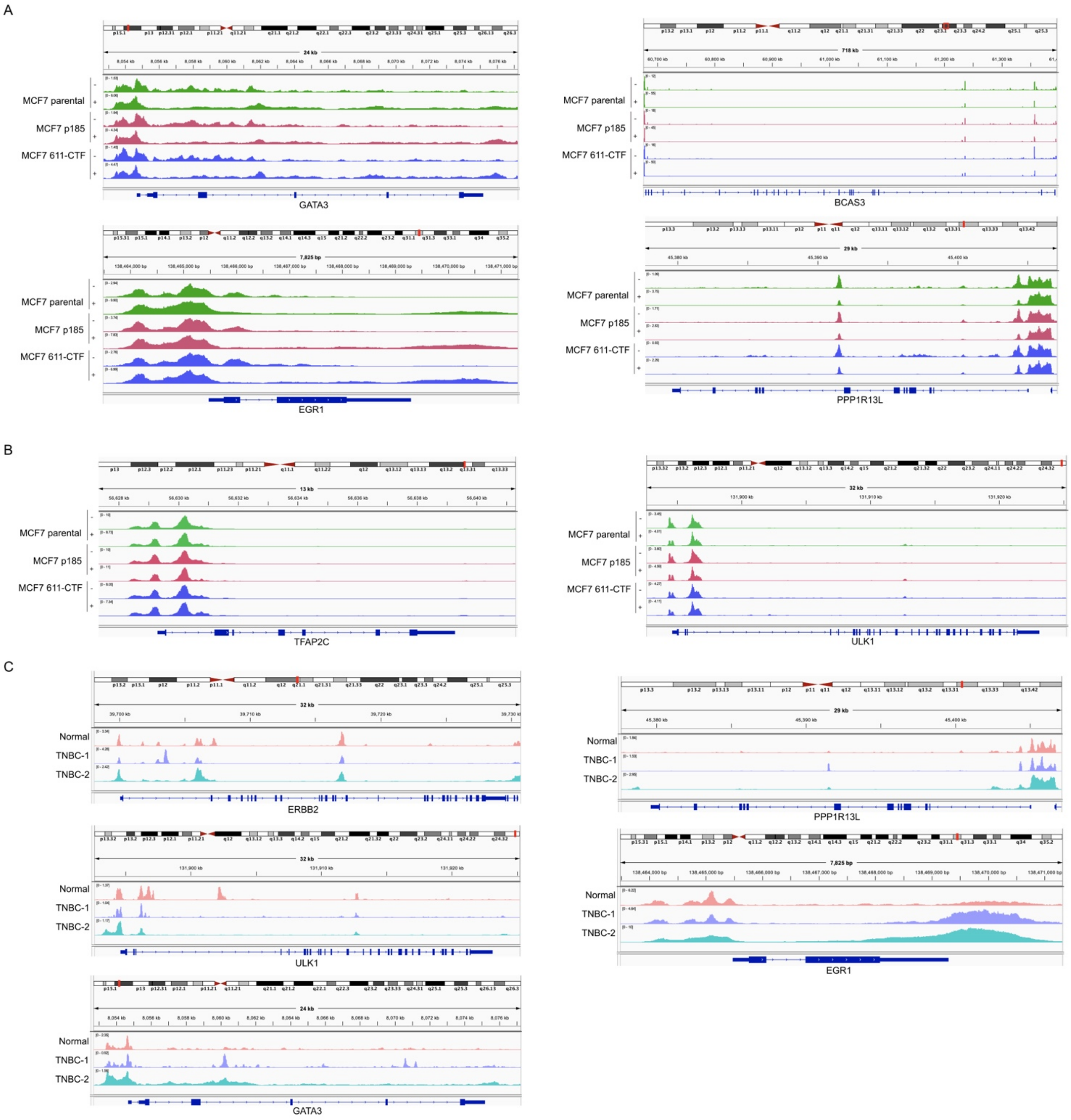

**Supplementary Figure 12: Protein analysis in transfected MCF7 cell lines upon lapatinib treatment.**

A) Location of the tested proteins on the volcano plot (DMSO control vs lapatinib treated). B) PCA plot of the proteins divided into treated and untreated conditions for each cell line. C) Bar blot with the selected tested proteins (ERBB2, ULK1, TFAP2C) on the 3 cell lines. Conditions with no expression were not plotted. The y-axis represents the protein expression level normalized by the expression of β-tubulin in the same cell. Proteins labeled with [Form 1 or 2] presented proteoforms in these gels. Form 1 corresponds to the observed proteoform with higher molecular weight, while Form 2 is the lower molecular weight form. Significant differences were shown as * (P<0.05).

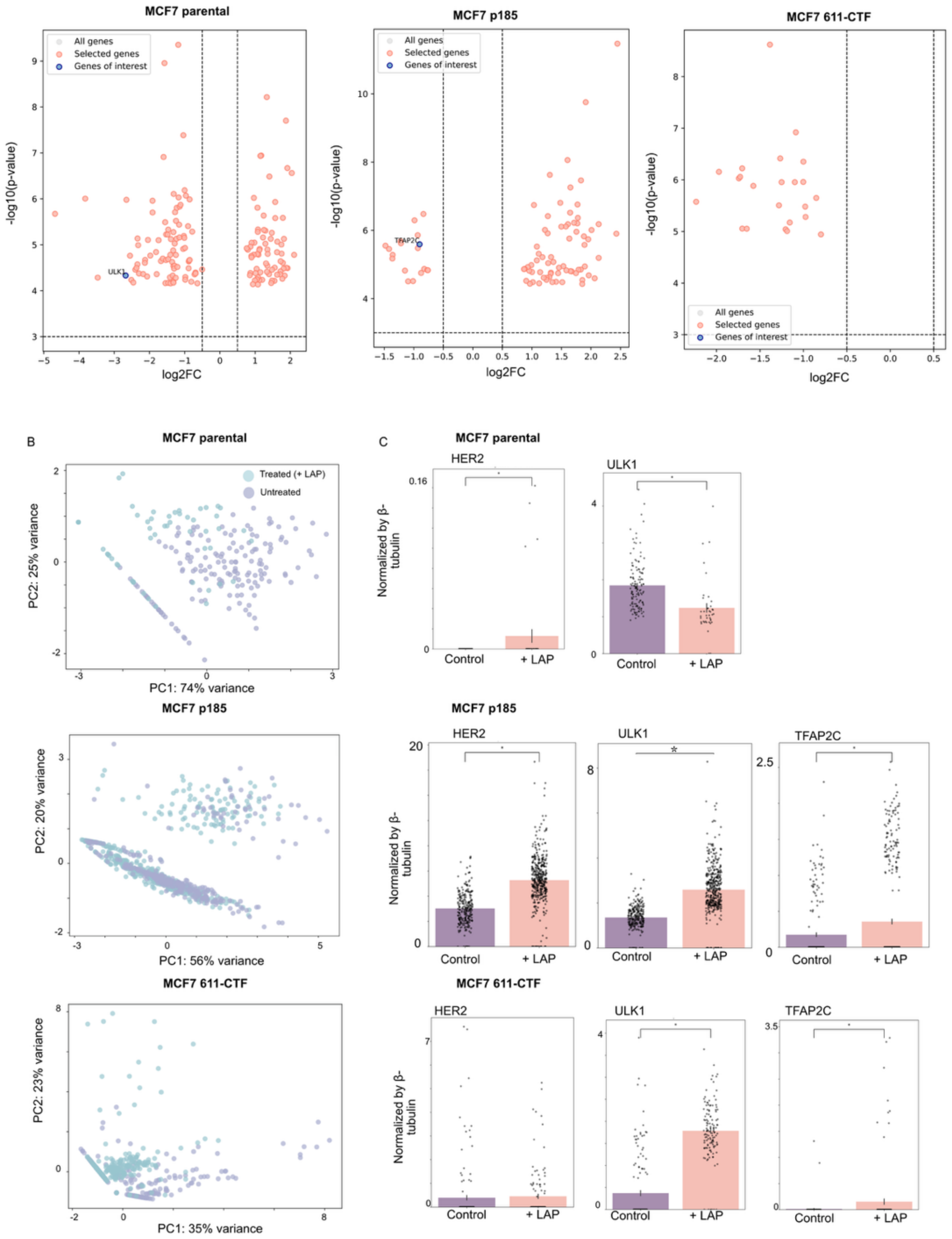

**Supplementary Figure 13: Immunofluorescence of tumor organoids.**

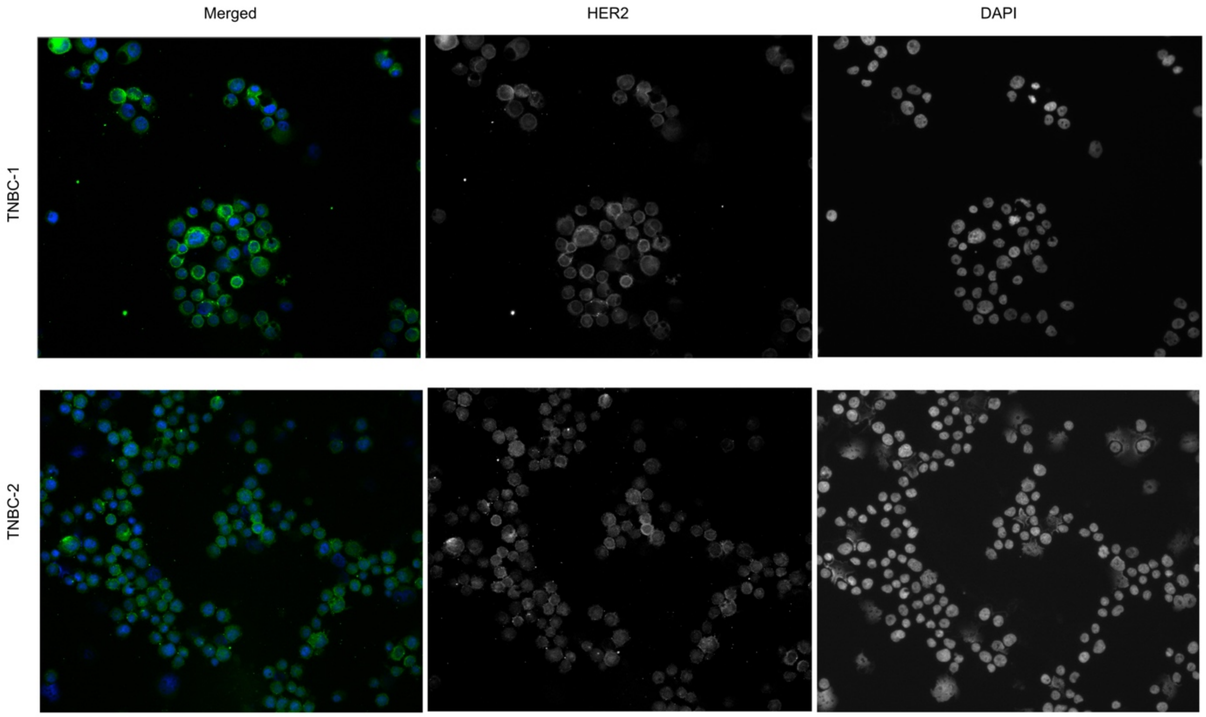

**Supplementary Figure 14: KEGG pathway analysis of organoids.**
A and B) KEGG analysis of the TNBC organoids, compared to the Normal organoid. C) Changes in chromatin accessibility of the *ERBB2* locus for Normal, TNBC-1 and TNBC-2.

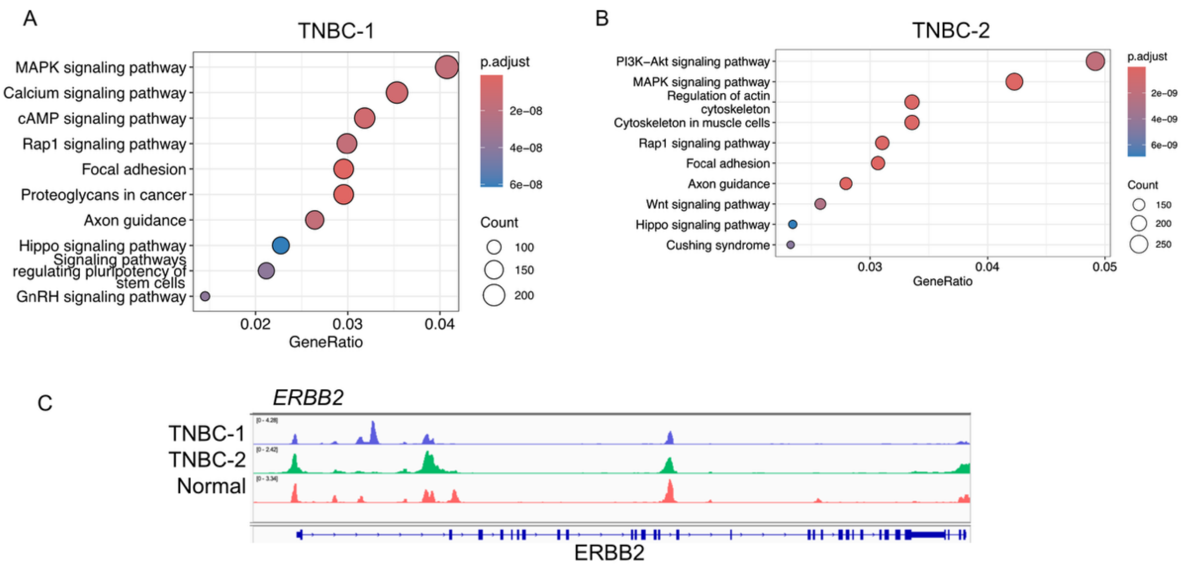

**Supplementary Figure 15: Protein expression of the organoids Normal, TNBC-1, and TNBC-2.**

A) Location of the tested protein on the volcano plot (TNBC vs Normal organoid). B) PCA plot of the proteins representing the three organoids. The star signs show the different subpopulation we could observe in TNBC-1, which expressed a HER2 p95 proteoform. C) Micrographs showing undetermined phenotypes (n=26), which were excluded from the analysis. These scWB bands displayed a very faint signal in the migration distance corresponding to p95HER2, but the signal intensity was insufficient for reliable classification. These micrographs are shown without background subtraction owning to the faint signal. D) Bar blot with the selected tested proteins (ERBB2, ULK1, GATA3, EGR1, PPP1R13L) on the 3 cell lines. Conditions with no expression were not plotted. The y-axis represents the protein expression level normalized by the expression of β-tubulin in the same cell. Proteins labeled with [Form 1 or 2] presented proteoforms in these gels. Form 1 corresponds to the observed proteoform with higher molecular weight, while Form 2 is the lower molecular weight form. Significant differences were shown as * (*P*<0.05).

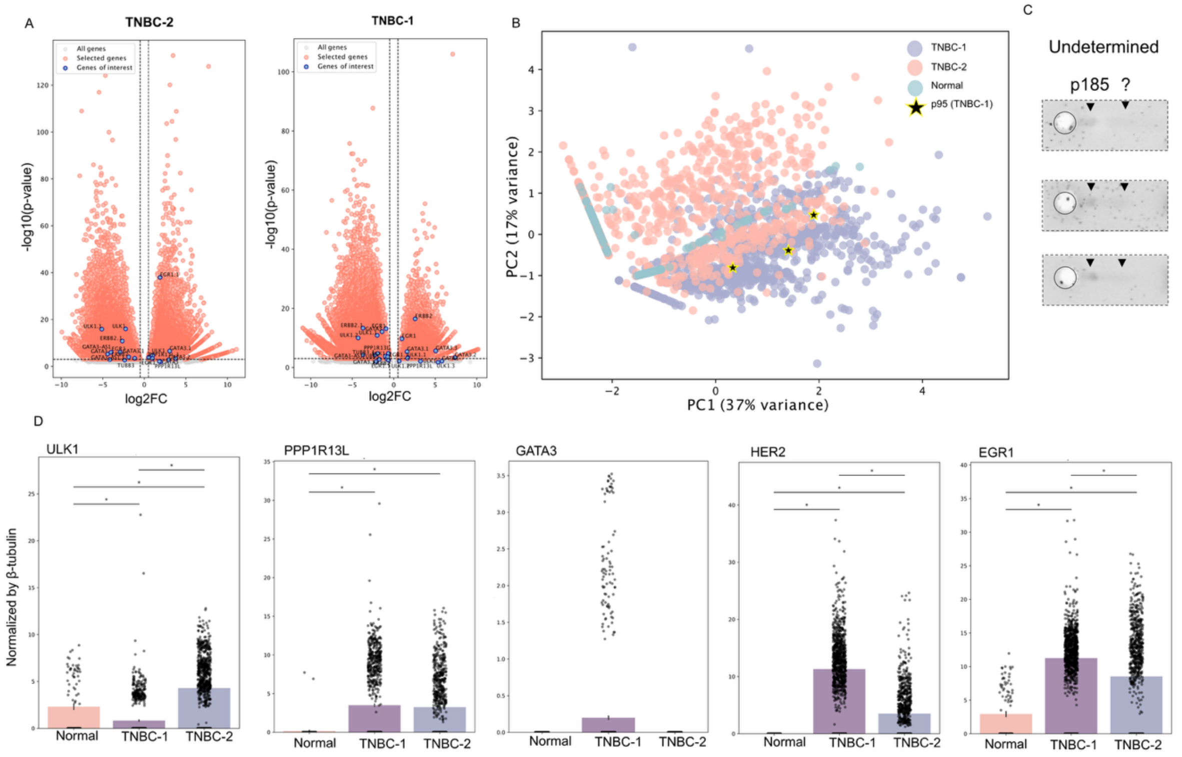

**Supplementary Table 1. Characteristics of the tested PDOs.**

| Organoid ID | Tissue type | Reference |
|---|---|---|
| Normal | Normal breast – BRCA1 mutation carrier | [8] |
| TNBC-1 | TNBC Grade 3 | [9] |
| TNBC-2 | TNBC Grade 2 | [10] |

Inflammation. Cancer Immunology Research *13*, 229–244. https://doi.org/10.1158/2326-6066.CIR-24-0416.
